# miR-7 controls glutamatergic transmission and neuronal connectivity in a Cdr1as-dependent manner

**DOI:** 10.1101/2023.01.26.525729

**Authors:** Cledi A. Cerda Jara, Seung Joon Kim, Gwendolin Thomas, Zohreh Farsi, Grygoriy Zolotarov, Elisabeth Georgii, Andrew Woehler, Monika Piwecka, Nikolaus Rajewsky

## Abstract

The circular RNA (circRNA) Cdr1as is conserved across mammals and highly expressed in neurons, where it directly interacts with microRNA miR-7. However, the biological function of this interaction is unknown. Here, using primary forebrain murine neurons, we demonstrate that stimulating neurons by sustained depolarization rapidly induced two-fold transcriptional up-regulation of Cdr1as and strong post-transcriptional stabilization of miR-7. Cdr1as loss caused doubling of glutamate release from stimulated synapses and increased frequency and duration of local neuronal bursts. Moreover, periodicity of neuronal networks was increased and synchronicity was impaired. Strikingly, these effects were reverted by sustained expression of miR-7 which also cleared Cdr1as molecules from neuronal projections. Consistently, without Cdr1as, transcriptomic changes caused by miR-7 overexpression were stronger (including miR-7-targets down-regulation) and enriched in secretion/synaptic plasticity pathways. Altogether, our results suggest that in forebrain neurons Cdr1as buffers miR-7 activity to control glutamatergic excitatory transmission and neuronal connectivity important for long-lasting synaptic adaptations.

## Introduction

For about ten years, it has been known that the circular RNA Cdr1as, originally discovered by the Kjems lab (Hansen et al., 2011), is one of the most highly expressed, vertebrate-conserved, circular RNAs in the mammalian brain (Memczak et al., 2013). Moreover, the expression of the microRNA miR-7 is deeply conserved, and is one of its most frequent direct targets (out of all coding or non-coding RNAs) in the mammalian brain is Cdr1as, which harbors more than hundreds of highly conserved miR-7 binding sites (Hansen et al., 2013; Memczak et al., 2013; Piwecka et al., 2017) and is specifically and highly expressed in excitatory neurons across the forebrain (Piwecka et al., 2017; Rybak-Wolf et al., 2015). But what is the function of this unusual (no other known circular RNA has this many conserved binding sites for a specific miRNA) and deeply conserved interaction between two non-coding RNAs?

miR-7 is an ancient bilaterian miRNA, evolutionarily conserved across vertebrates and considered to be a prototypical neuroendocrine miRNA (Kredo-Russo et al., 2012; Latreille et al., 2014). Extensive comparative data indicate that miR-7 evolved within the neurosecretory brain (Christodoulou et al., 2010), and that the neuronal and pancreatic differentiation lineages are closely related (Zhao et al. 2007). Some roles of miR-7 in mammalian forebrain development have been described, mainly by its interaction with target gene Pax6 in neuronal progenitors (Pollock et al., 2014; Zhang et al., 2018), but nothing is known about miR-7 role in post-mitotic forebrain. Mature miR-7 is enriched in central nervous system neuroendocrine glands and pancreatic tissues (Bravo-Egana et al., 2008; Hsu et al., 2008; Landgraf et al., 2007). In mouse pancreas, miR-7 has been shown to function as a negative regulator of stimulus-dependent insulin secretion (Latreille et al., 2014; Xu et al., 2015). This mechanism seems to be conserved for insulin-producing cells across the animal kingdom (Agbu et al., 2019; Latreille et al., 2014). In view of these findings, we hypothesized that miR-7 might regulate the release of key neuronal transmitters and that Cdr1as acts to control miR-7 function in forebrain neurons.

Because some circRNAs are induced in neurites by homeostatic plasticity stimulation (You et al., 2015), we also speculated that the Cdr1as-miR-7 system may function mainly in stimulated neurons – moreover, (as we show), in unstimulated forebrain neurons miR-7 is only lowly expressed.

To test these ideas, we performed sustained neuronal stimulations and sustained perturbation of miR-7 expression in Cdr1as WT and KO primary forebrain neurons to investigate the dynamic interplay between Cdr1as and miR-7 in excitatory neurotransmission and neuronal network activity (synaptic plasticity). We show that Cdr1as acts as a buffer that controls dynamic miR-7 regulation of glutamate release from presynaptic terminals, and that this regulation strongly modulates neuronal connectivity responsible for long-lasting adaptation of synaptic plasticity.

## Results

### Sustained neuronal depolarization transcriptionally induces Cdr1as and post-transcriptionally stabilizes miR-7

To test our hypothesis that miR-7 is involved in forebrain neurotransmitter secretion in similar ways as it is participating in stimulus-regulated secretion in pancreatic cells and neurosecretory glands (LaPierre et al., 2022; Latreille et al., 2014) and because some circRNAs were shown to be modulated specifically by neuronal activity (You et al., 2015), we tested the response of Cdr1as and miR-7 to sustained neuronal depolarization in primary forebrain neurons from mice (**Fig. 1A**). Elevated extracellular K^+^ generates a global neuronal depolarization. Therefore, extracellular KCl treatments are useful for elucidating signalling, transcriptional, and plasticity-related events associated with homeostasis, and long-term potentiation, which controls which neurons are recruited for sensory stimuli, learning, memory, etc. (Rienecker et al., 2020). A fast, primary response of neuronal stimulation are activity-regulated immediate early genes (IEGs), dependent on the duration of the stimulation (Bartel et al., 1989; Tyssowski et al., 2018). More precisely, IEG are well known to be the primary transcriptional, rapid, and transient response to neuronal activation and depolarization stimuli and these transcripts encode mostly transcription factors and DNA-binding proteins, (Curran & Morgan, 1987; Herdegen & Leah, 1998; Lanahan & Worley, 1998; Tischmeyer & Grimm, 1999).

**Figure 1.**
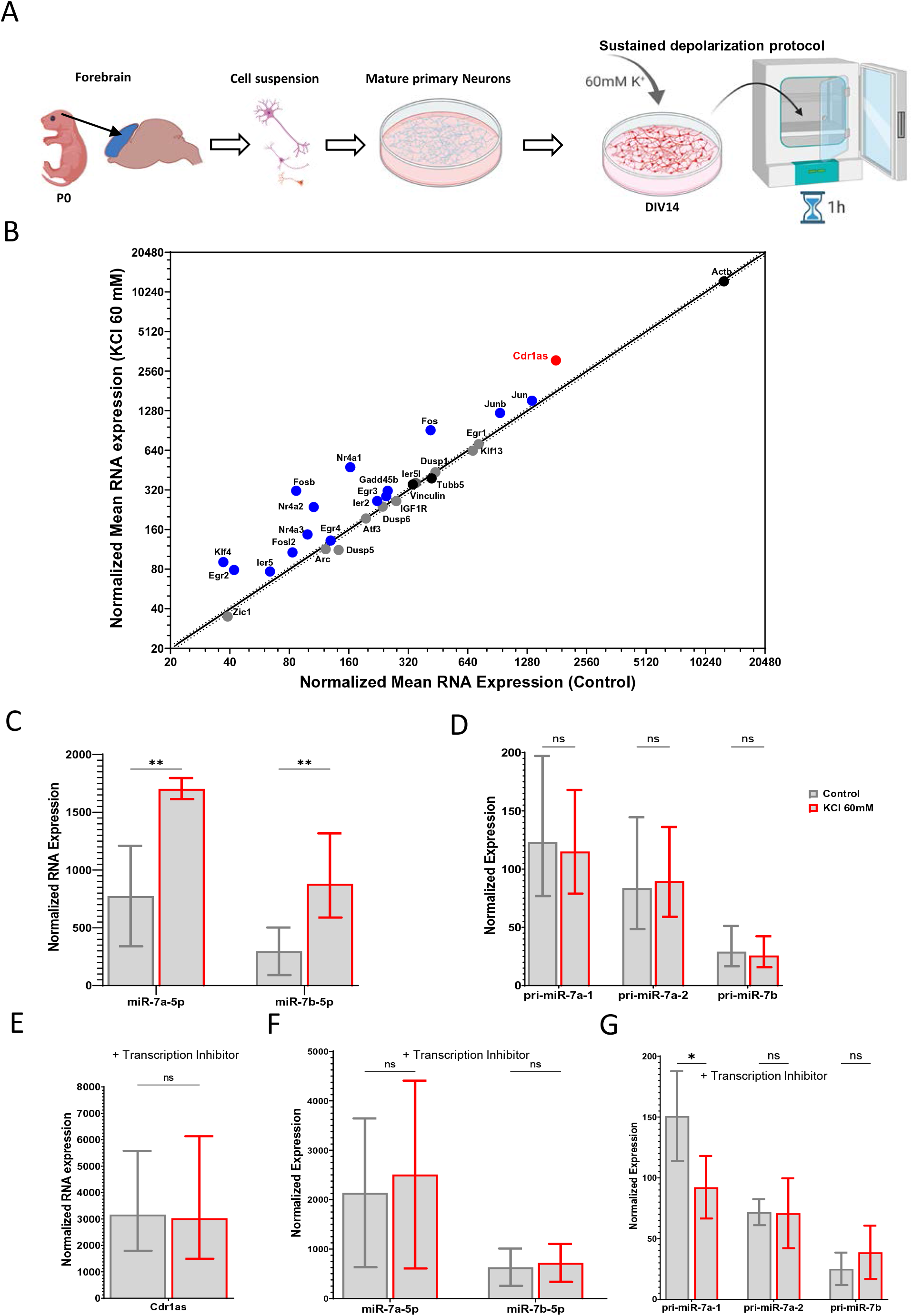
Sustained neuronal depolarization causes transcriptional up-regulation of Cdr1as and post-transcriptional stabilization of miR-7. **(A)** Protocol to culture primary forebrain neurons (modified from Keach and Banker, 2006:Methods). DIV14 neurons, pretreated with TTX, CNQX and AP5, were stimulated with 1 hour of KCl (60mM). **(b)** RNA quantification before and after KCl treatment (Nanostring nCounter, Methods). Statistically significantly upregulated RNA levels are shown in blue (IEGs) and in red (Cdr1as). Housekeeping genes: black. RNA counts are normalized to housekeeping genes (*Actb, Tubb5 and Vinculin*). Each dot represents the mean of 4 biological replicates (4 independent primary cultures from 4 animals). Black line: linear regression; dotted line: 95% confidence interval. **(C)** Expression levels of mature miR-7 isoforms quantified by small RNA-seq for 3 independent primary cultures, before and after sustained depolarization. Bar plot: the mean. P value: ratio paired t-test (significance: p value < 0.002). Error: SD **(D)** Quantification of primary miRNAs (Nanostring nCounter, Methods), before and after sustained depolarization. RNA counts are normalized to housekeeping genes (Actb, Tubb5 and Vinculin). Bar plot represents the mean of 4 biological replicates (4 independent primary cultures from 4 animals). P value: ratio paired t-test. Error: SD. **(E)** Quantification of Cdr1as (Nanostring nCounter, Methods), before and after sustained depolarization plus pre-incubation with transcription inhibitor (DRB). RNA counts are normalized to housekeeping genes (*Actb, Tubb5 and Vinculin*), bar plot represents the mean of 3 biological replicates (3 independent primary cultures from 3 animals). P value: ratio paired t-test. Error: SD. **(F)** Expression levels of mature miR-7 isoforms quantified by small RNA-seq for 3 independent primary cultures, before and after sustained depolarization plus pre-incubation with transcription inhibitor (DRB), bar plot represents the mean. P value: ratio paired t-test. Error: SD **(G)** Quantification of primary miRNAs (Nanostring nCounter, Methods),before and after sustained depolarization plus pre-incubation with transcription inhibitor (DRB). RNA counts are normalized to housekeeping genes (*Actb, Tubb5 and Vinculin*). Bar plot represents the mean of 3 biological replicates (3 independent primary cultures from 3 animals). P value: ratio paired t-test. Error: SD.

We pre-treated primary neurons at DIV13 with TTX, CNQX and AP5 overnight to inhibit spontaneous neuronal responses generated by Na^+^-dependent depolarization or glutamate release. On DIV14, the culture media was exchanged with media containing high extracellular concentrations of potassium (KCl 60 mM). This treatment was performed for 1 hour (control: media with the same osmolarity as the high potassium media) (**Fig. 1A, right**).

The observed consistent and highly significant up-regulation of IEG in K^+^ treated neurons (including master neuronal activity regulators such as *Fos, Fosb, Jun, Junb, Egr2, Nr4a1, Nr4a2*, **Fig. 1B blue dots**) is consistent with previously described gene expression changes for one hour of sustained K^+^ treatment (Bartel et al., 1989, p. 89; Rienecker et al., 2020). Genes not expected to react to stimulation were indeed not changing in expression, for example house-keeping genes (*Actb, Vinculin, Tubb5*), other transcription factors (*Klf13, Atf3, Zic1*) and cell-cycle related genes (*Dusp1, Dusp5, Dusp6, IGF1R*) (**Fig. 1B, black and grey dots**).

Together with higher expression of activity-dependent IEGs, we observed a robust and significant up-regulation of Cdr1as similar in strength to what we have seen for IEGs (**Fig. 1B**, **red label**). In fact, the absolute expression levels of Cdr1as were the highest observed across all genes tested. Moreover, of all circRNAs that we detected by total RNA sequencing (Methods), Cdr1as was by far the most highly expressed and induced circRNA (**Suppl. Fig. 1J**). Mature miR-7 isoforms (miR-7a and miR-7b), were also strongly up-regulated after K+ stimulation, compared to the control (**Fig. 1C**). No changes were observed in the expression of mature miR-671, involved in Cdr1as turnover, or in highly expressed neuronal miRNA Let-7a (**Suppl. Fig. 1B**).

Interestingly, none of the primary transcripts of miR-7 (pri-miR-7a-1, pri-miR-7a-2, pri-miR-7b) changed after K+ stimulation (**Fig. 1D**). This shows that the regulation of mature miR-7 occurred post-transcriptionally. We further probed the dependence on transcription by blocking transcription with an elongation inhibitor (DRB) before and without K+ treatment (**Fig. 1E-F-G**). For none of the genes significant up-regulation was observed. Down-regulation was observed for some IEGs and primary miR-7 isoforms, probably due to their short half-life times (**Suppl. Fig. 1C-E and Fig. 1G**). Cdr1as levels were unperturbed (**Fig. 1E**), as well as mature miR-7 levels were unchanged, likely explained by the known high stability of Cdr1as (Memczak et al., 2013) and the known stability of miR-7 bound to AGO (Bartel, 2018).

We also tested the response of the Cdr1as RNA network to the same sustained depolarization in primary astrocytes exposed to the neuronal culture (glial feeder layer) and we did not observe any response after K^+^ stimulation, with or without DRB pre-treatment, compared to the control. Interestingly, we also did not observe any up-regulation of IEGs, Cdr1as, Cyrano, pri- or mature miR-7 or of any other miRNAs tested (**Suppl. Fig. 1F-H**), suggesting that Cdr1as and mir-7 activity-dependent regulation is part of a neuronal-specific response mechanism.

Together our observations demonstrate that miR-7 is post-transcriptionally regulated in a likely neuronal-specific way, either by regulation of maturation of the miR-7 precursor or by stabilization via interaction with Cdr1as. Strikingly, while most IEGs up-regulated by depolarization are transcription factors encoding nuclear proteins, Cdr1as is enriched in the neuronal cytoplasm (**Suppl. Fig. 1I**) and broadly expressed in neurites (**Suppl. Fig. 2A**), where Cdr1as binds mature miR-7 (Hansen et al., 2013; Kleaveland et al., 2018; Memczak et al., 2013; Piwecka et al., 2017). Considering this difference, together with the neuronal stimulation observations, we hypothesize that the cellular role of Cdr1as in neurons might be to safeguard or buffer the action of miR-7 in stimulated neurons.

### Synaptic terminals of Cdr1as-KO neurons show strongly increased presynaptic glutamate release after stimulation

To test if Cdr1as is involved in synaptic transmission in stimulated neurons, we used the forebrain primary neurons of Cdr1as-KO animals (Piwecka et al., 2017) with their corresponding WT littermates. We performed RNA quantification of marker genes at DIV21 and did not observe significant differences in the expression of neuronal (excitatory and inhibitory), glial, or proliferation and maturation RNA markers between cultured Cdr1as-KO and WT neurons (**Suppl. Fig. 2C**). In the forebrain, glutamatergic neurons are the main cell type (Niciu et al., 2012), and we corroborated that this is also true in our primary neuronal cultures from WT as well as Cdr1as-KO animals, glutamatergic neurons were the predominant cell type (**Suppl. Fig. 2C**). We then focused on measuring the activity of glutamate, as the main neurotransmitter relevant in our primary cultures.

We used a genetically-encoded glutamate sensor (GluSnFR (Marvin et al., 2013)) to perform real-time recordings of spontaneous and action potential-evoked (APs; electrical stimulation) glutamate release from individual presynaptic terminals in mature neurons DIV18 to 21, pre-treated with TTX, CNQX and AP5, to inhibit spontaneous or post-synaptic responses. Then, separately, we calculated the frequency of spontaneous neurotransmitter release over a time (5 minutes of analysis) and the probability of evoked glutamate release in individual synapses during a sustained trend of electrical stimulations (20 APs at 0.5 Hz), as previously described by Farsi et al., 2021 (**Fig. 2A and Suppl. Fig. 2D**).

**Figure 2.**
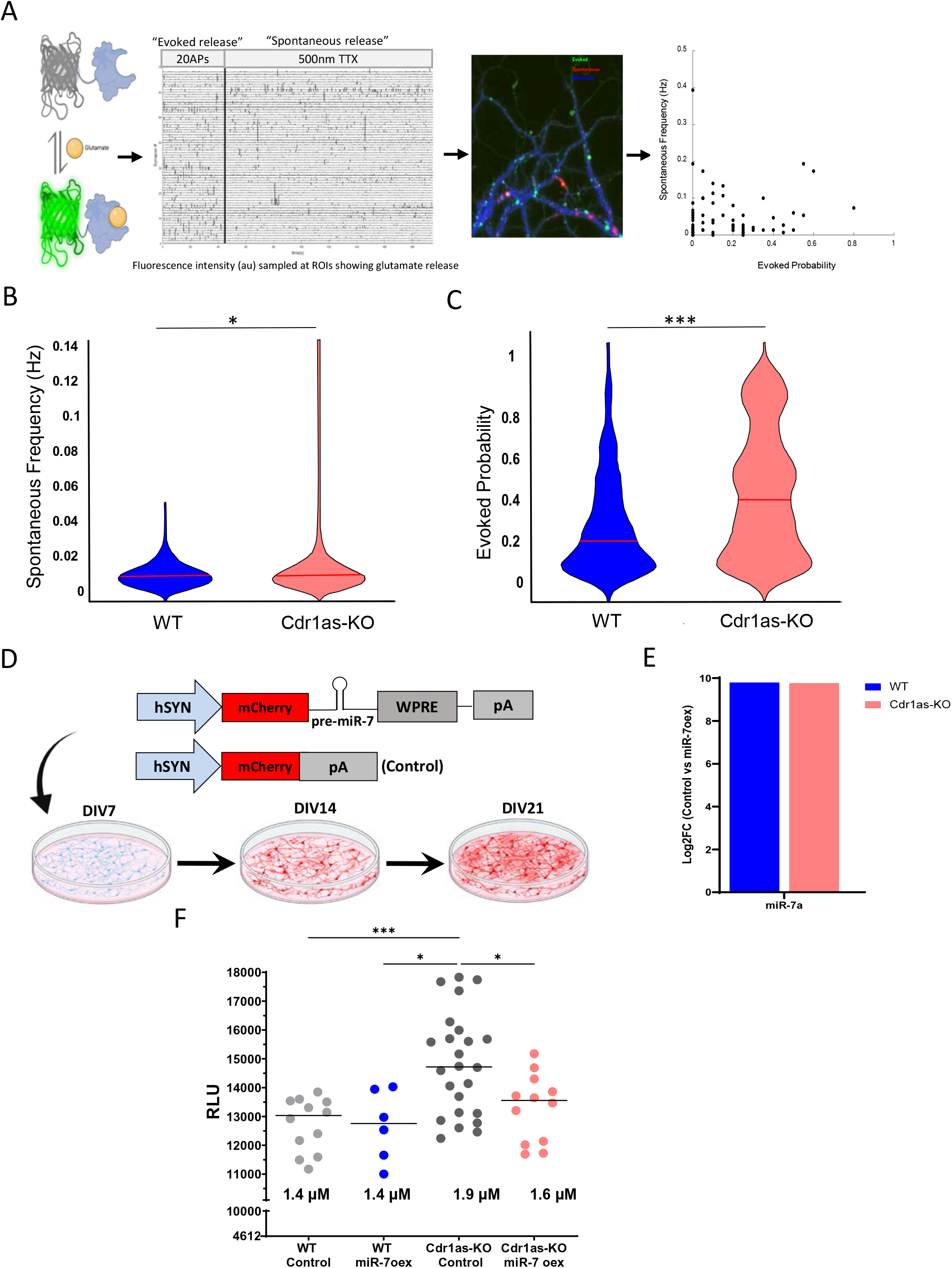
Presynaptic terminals of Cdr1as-KO forebrain neurons show increased glutamate release which is reverted by sustained miR-7 overexpression. **(A)** Transduction of a glutamate sensor (AAV, Methods) into WT and Cdr1as-KO primary neurons followed by real-time imaging of excitatory synaptic terminals during AP-evoked (20 APs at 0.5 Hz) and spontaneous (5 minutes + 500 nM TTX) release conditions. **(B)** Quantification of spontaneous glutamate release calculated as spontaneous frequency. Violin plots: integration of the synapses from all animals used (WT: n = 4 animals and 662 synapses; Cdr1as-KO: n = 23 animals and 421 synapses; Methods).Synapses with fluorescence value = 0 were removed. Kernel-Density-Estimate plotted (Methods). Red line: median. P value: Unpaired t-test. **(C)** Quantification of AP-evoked glutamate release calculated as evoked probability. Violin plots: integration of the synapses from all animals used (WT: n = 4 animals and 335 synapses; Cdr1as-KO: n = 3 animals and 237 synapses; Methods).Synapses with fluorescence value = 0 were removed. Kernel-Density-Estimate plotted (Methods). Red line: median. P value: Unpaired t-test. **(D)** Schematic representation of the miR-7a overexpression and the mCherry control constructs. Viral transduction performed at DIV7 (AAV, Methods). mCherry became visible at DIV14 and experiments were performed at DIV21. **(E)** Quantification of miR-7a up-regulation by RNA-Seq (Methods) in WT and Cdr1as-KO neurons at DIV21, 14 days post miR-7a overexpression. Bar plots represents mean of 4 independent biological replicates per genotype. **(F)** Glutamate secretion assay performed on neuronal media collected from DIV18-21 WT and Cdr1as-KO neurons infected with control or miR-7a overexpression construct. Each dot represents the glutamate concentration quantified in an independent primary culture (WT = 12; WT + miR-7 overexpression = 12; Cdr1as-KO = 12; Cdr1as-KO + mirR-7 overexpression = 24 independent replicates). Glutamate concentrations are calculated based on a glutamate titration curve (Methods, Supplementary material). Baseline concentration of glutamate in control media without cells was 0,28 uM (4612 RLU). Black line: median. P value: two-way ANOVA, all other comparisons were not statistically significant (not shown).

Our data showed a mild but significant increase in spontaneous glutamate release in Cdr1as-KO presynaptic terminals as compared to WT neurons (**Fig. 2B**), consistent with previously shown up-regulation of spontaneous miniature EPSC frequency in Cdr1as-KO hippocampal patch clamp experiments of single, isolated neurons (autapses, Piwecka et al., 2017). However, we observed that the increase in glutamate release was mainly driven by the spontaneous activation of some groups of synapses, rather than by a global activation of all terminals (**Fig. 2B, pink Kernel density distribution**). Strikingly, we observed much stronger up-regulation of action potential-evoked glutamate release in Cdr1as-KO neurons: 20 APs at 0.5 Hz for 40 secs. (**Fig. 2C**), suggesting that Cdr1as plays a potentially important role in the modulation of excitatory transmission during sustained stimulation, in line with our hypothesis derived from K^+^ stimulation of WT neurons (**Fig. 1**).

Overall, real-time recording of presynaptic neurotransmitter release revealed a direct link between Cdr1as dependent regulation of glutamatergic transmission in resting and particularly in stimulated neuronal states. How could Cdr1as regulate glutamatergic transmission? It has been proven that Cdr1as’ main binding partner, miR-7, directly acts as a negative regulator of stimulus-regulated secretion in secretory glands, where the miR-7 is within the top highly expressed miRNA (Bravo-Egana et al., 2008, p. 7; Landgraf et al., 2007; Latreille et al., 2014). However, forebrain neurons are characterized by a very low basal expression of mature miR-7 when compared to characteristic neuronal miRNAs (**Suppl. Fig. 2E and 3B**). Therefore, to reliably study the functions of miR-7 in forebrain neurons and to molecularly mimic the increased miR-7 levels observed as consequence of neuronal stimulation (K^+^ treatments; **Fig. 1**), we decided to induce miR-7 expression in un-stimulated WT and Cdr1as-KO neurons.

### Sustained miR-7 overexpression reverts increased glutamate secretion in Cd1as-KO neurons

We created a neuron-specific (human synapsin promoter; hSYN) long-term transgene overexpression system, consisting of a miR-7 precursor transcript (pre-miR-7a-1) coupled to a fluorescent reporter (mCherry) and packaged into an adeno-associated virus particle (AAV) for highly efficient neuron transduction, together with a fluorescent vector control) (**Fig. 2D**, Methods).

We infected WT and Cdr1as-KO primary neuronal cultures on DIV7 with miR-7 or control AAV particles and then monitored mCherry as a reporter of miR-7 expression, until DIV21 when neurons are synaptically mature (**Fig. 2D**). We then tested miR-7 overexpression and confirmed significant induction of mature miR-7 in WT and Cdr1as-KO cultures (**Fig. 2E**). Importantly, induced miR-7 levels were very similar in both cases (**Fig. 2E**). Additionally, we did not observe significant change of miR-671 or for the highly expressed neuronal miRNA let-7a in any genotype (**Suppl. Fig. 3C**), indicating that our overexpression did not interfere with the abundance of other miRNAs.

We observed that the mCherry protein became visible starting from DIV14 onwards (7dpi). Using single molecule RNA *in situ* hybridization of miR-7, we demonstrate that the overexpressed miRNA is found homogenously distributed in somas and neurites of neurons from both genotypes (**Suppl. Fig. 3D**), while in control neurons (not infected or empty control) the signal is sparse and only few molecules are captured (~40 copies per cell; **Suppl. Fig. 3A**). Accordingly, we kept DIV14 (7dpi) as the starting time-point to monitor neuronal activity in all following experiments.

We then measured secreted glutamate in the media of DIV18-21 WT and Cdr1as-KO neuronal cultures at resting state, in neurons infected with control or miR-7 overexpression AAV (14dpi). For this, we set up a bioluminescent enzymatic assay (Methods), based on the activity of glutamate dehydrogenase (GDH) coupled to the activity of Luciferase (Ultra-GloTM rLuciferasa, Promega), which results in the production of light proportional to the concentration of glutamate in the media (**Suppl. Fig. 3E**). Exact concentrations of secreted glutamate were calculated by interpolation of luminescence values (RLU) from a glutamate standard curve (**Suppl. Fig. 3F**).

Secreted glutamate levels were not changed in WT cells when overexpression miR-7 (**Figure 2F**), in contrast to other systems such as pancreatic beta cells where specific overexpression of miR-7 induced a reduction of insulin secretion (Latreille et al., 2014). However, consistent with our previous observations (**Fig. 2B, C**), glutamate levels were strongly and significantly increased in Cdr1as-KO neurons. Strikingly, this increase was reverted to almost WT levels when miR-7 was overexpressed in the absence of Cdr1as. As Cdr1as is absent or only spuriously expressed in the pancreas, our data suggest that Cdr1as in forebrain neurons has evolved to buffer miR-7 function.

### Sustained miR-7 overexpression reverted increased local neuronal bursting in Cdr1as-KO neurons

To investigate more detailed neuronal activity differences associated with the loss of Cdr1as, along with examining if and how those differences are associated with miR-7, we recorded local and network real-time activity over neuronal maturation (DIV7 until DIV21), in WT and Cdr1as-KO cultures, with or without miR-7 overexpression. We used a multi-electrode array well-based system (MEA, Axion BioSystems) to record extracellular field APs (neuronal spikes), and ultimately network activity (**Fig. 3A**).

**Figure 3.**
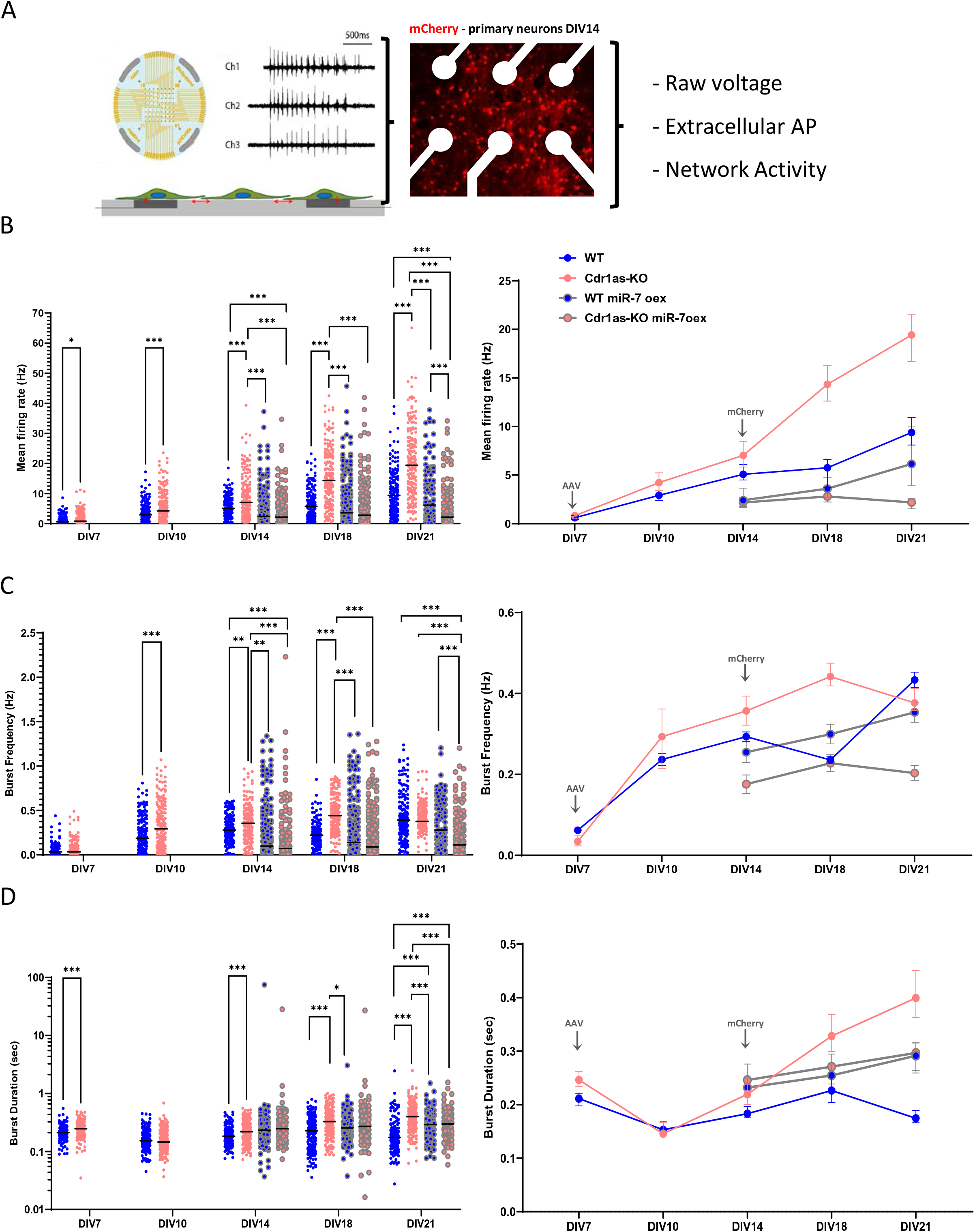
Removal of Cdr1as causes increased frequency and duration of local action potentials and neuronal bursting. Sustained miR-7 expression reverts these effects. **(A)** Scheme of the Multi electrode Array recording protocol (Axion Biosystems, CytoView MEA 48, Methods). Second panel: representative image of cultured neurons DIV14 in a recording well. white: electrodes, red: mCherry reporter. Third panel: schematic representation of output data, extracellular field potentials and neuronal network activity. AP: Adaptative threshold 6 SD. Sampling frequency 12.5 kHz. Active electrode selection criteria 5 spikes/minute. **(B)** Mean Firing Rate: Total number of spikes per single-electrode divided by the duration of the analysis (600s), in Hz. Left panel: each dot represents a single electrode recording from 4 independent primary cultures (WT = 187; WT + miR-7 overexpression = 194; Cdr1as-KO = 189; Cdr1as-KO + miR-7 overexpression = 200 electrodes). Horizontal line: median. Right panel: median value across timepoints with 95% confidence interval is plotted; arrows indicate transduction time (AAV, DIV7) and reporter first visualization (mCherry, DIV14), respectively. P value: two-way ANOVA, all other comparisons were not statistically significant (not shown). **(C)** Burst Frequency: Total number of single-electrode bursts divided by the duration of the analysis, in Hz. Left and right panels plotted as in (B). **(D)** Burst Duration: Average time (sec) from the first spike to last spike in a single-electrode burst. Left and right panels plotted as in (B).

When measuring local APs recorded by an electrode placed nearby the generating neurons, we observed that throughout maturation time (and similar to our data for glutamate secretion in media), there are no significant changes in the frequency of firing in WT neurons overexpressing miR-7 as compared to WT controls in any of the analyzed timepoints (**Fig. 3B, blue and grey-blue datapoints**). However, Cdr1as-KO neurons consistently showed a significant up-regulated spontaneous firing frequency (larger number of functional APs per second), which progressively increased from DIV7 until DIV21 as compared to WT neurons (**Fig. 3B, pink and blue datapoints**). Again, this up-regulation was reverted to almost WT levels (or exceeding those) when over-expressing miR-7 in Cdr1as-KO neurons (**Fig. 3B pink and grey-pink datapoints**). This “rescue” was the strongest in more mature neurons (DIV21). Together, we conclude that our observations of up-regulated glutamate secretion in Cdr1as-KO (**Fig. 2**) translate into increased functional APs.

To test if changes in firing rates are linked to changes in patterns of neuronal activity, we measure neuronal burst formation (defined as rapid AP spiking followed by inactive periods much longer than typical inter-spike intervals; Methods). In WT neurons no significant differences in bursting frequency were observed after sustained overexpression of miR-7 compared to controls, at any timepoint (**Fig. 3C, blue and grey-blue**). However, Cdr1as-KO neurons bursting activation patterns were persistently increased over maturation. This increase in bursting was as well reverted by miR-7 overexpression to WT or even lower frequencies (**Fig. 3C, pink and grey-pink datapoints**).

Cdr1as-KO neurons also showed significantly longer burst durations, (**Fig. 3D, pink and blue datapoints**) and a higher number of functional spikes per burst (**Suppl. Fig. 4A and 4B, pink and blue datapoints**), both growing in deviation from WT as the neurons matured. Interestingly, sustained miR-7 overexpression only significantly affected burst duration in DIV21 neurons and it did so in opposite ways, depending on the neuronal genotype, increasing the duration in WT neurons, and decreasing it in Cdr1as-KO.

**Figure 4.**
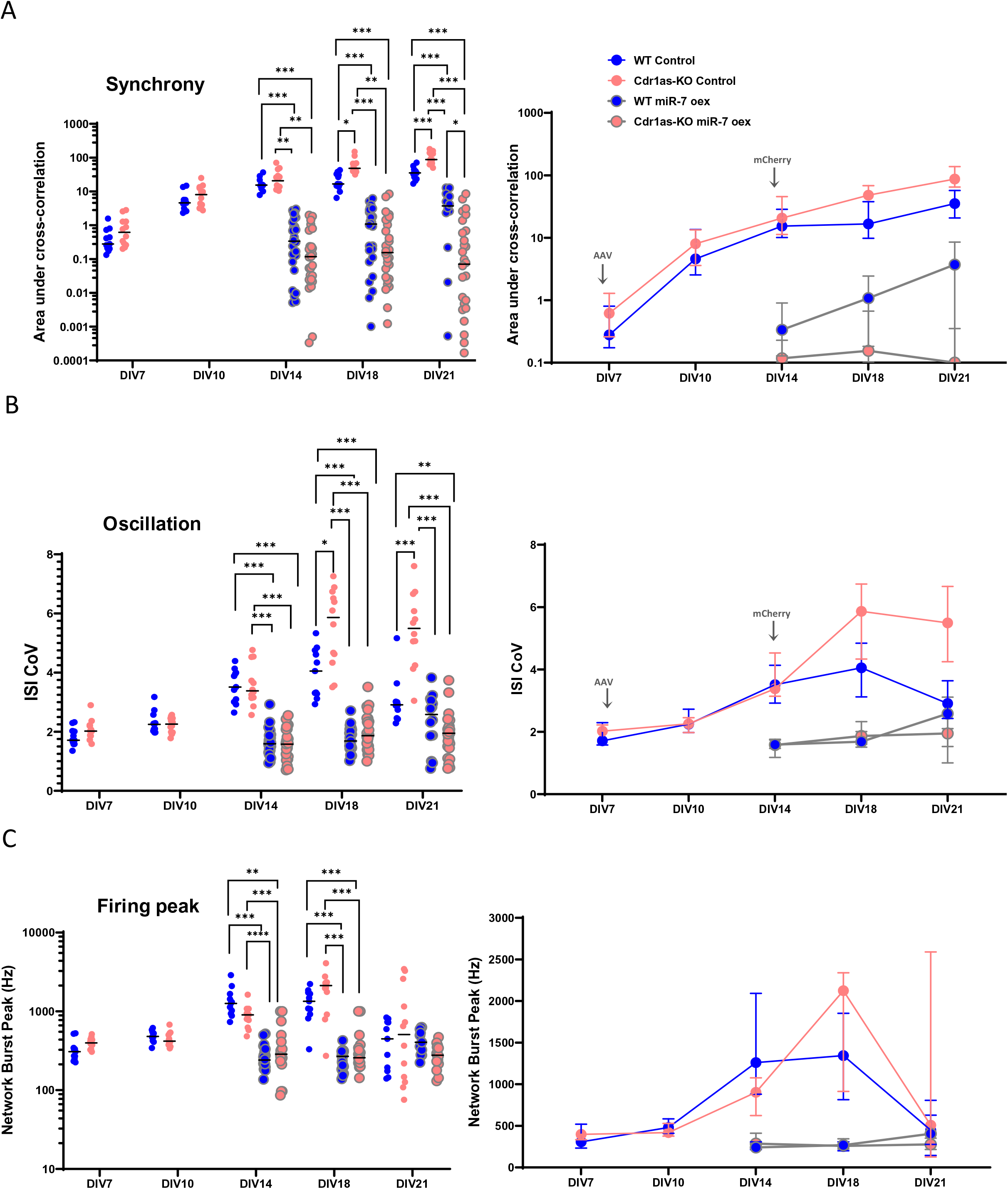
Removal of Cdr1as affects neuronal connectivity by changing network synchrony and oscillation. Sustained miR-7 expression reverted these effects. **(A)** Asynchrony: The ability of neurons to generate APs simultaneously was calculated as area under the well-wide pooled inter-electrode cross-correlation. Higher areas indicate lower synchrony (Halliday et al., 2006, Methods). Left panel: each dot represents a network recording (Methods) from 4 independent primary cultures (WT = 11; WT + miR-7 overexpression = 26; Cdr1as-KO = 12; Cdr1as-KO + miR-7 overexpression = 28 electrodes). Horizontal line: median. Right panel: median across timepoints with 95% CI plotted, arrows indicate transduction time (AAV, DIV7) and reporter first visualization (mCherry, DIV14). All metrics apply to network bursts across single wells within 20 ms. P value: two-way ANOVA, all other comparisons were not statistically significant (not shown). **(B)** Oscillation: Average across network bursts of the inter spike interval coefficient of variation (ISI CoV) (standard deviation/mean of the inter-spike interval) within network bursts. Oscillation is a measure of how the spikes from all of the neurons are organized in time. WT = 11-26; Cdr1as-KO: 12-28 independent network recordings. Left and right panels plotted as in (A). **(C)**Burst Peak: Maximum number of spikes per second in the average network burst. The peak of the average network burst histogram divided by the histogram bin size to yield spikes per sec (Hz). WT = 11-26; Cdr1as-KO: 12-28 independent network recordings. Left and right panels plotted as in (A).

Altogether, the real-time recording of local neuronal activity not only demonstrated that AP frequencies are impaired in Cdr1as-KO neurons, but suggested that cellular mechanisms responsible for transforming local AP into coordinated activation patterns (bursting activity) are affected in Cdr1as-KO neurons and all defects are effectively rescued by sustained mir-7 overexpression. Longer, more frequent, higher number of spikes per bursts are indications of stronger excitatory transmission, or of less inhibition, as it takes longer time to shut down a burst (Zeldenrust et al., 2018), therefore, our local neuronal activity results suggest that the effect of the interaction of Cdr1as and miR-7 on glutamatergic release (**Fig. 2**) is linked to the modulation of some aspects of synaptic plasticity, neural coding, and network synchronization.

### Sustained miR-7a up-regulation strongly affects neuronal connectivity in a Cdr1as dependent manner

To investigate further how Cdr1as and miR-7 are relevant to the modulation of synaptic plasticity and neuronal network patterns, we explored more complex neuronal activity parameters, taking advantage of multi-electrode simultaneous measurements.

First, we measured network synchrony, by comparing the activity profiles of two neighboring electrodes across different time delays (area under the cross correlation). When the area is 0 there is perfect synchrony, while larger area values indicate lower synchronicity. We observed a more asynchronous network in Cdr1as-KO neurons at late states of maturation (DIV14 to DIV21) when compared to WT (**Fig. 4A and Suppl. Fig. 4C, pink and blue datapoints**), and after sustained miR-7 overexpression the synchronicity was significantly increased for both genotypes. The size of this miR-7 dependent effect was much stronger in the case of Cdr1as-KO neurons (**Fig. 4A, grey-blue and grey-pink datapoints**). This higher synchrony could also be explained by less neuronal activity after miR-7 overexpression, as shown in our local neuronal activity observations (**Fig. 3**).

Second, we analyzed AP oscillations across the network and captured the distribution of action potentials. We measured the coefficient of variation of the intervals inter-spike (ISI CoV), where zero (0) values indicate perfect Poisson distribution of APs and higher values represent irregular bursting. We observed significantly higher oscillation periodicity in Cdr1as-KO network at very late points of maturation (DIV18-DIV21). In WT and Cdr1as-KO neurons, after sustained miR-7 overexpression there was a significant down-regulation of oscillatory bursting (**Fig. 4B, grey-blue and grey-pink datapoints**). The size of the miR-7 dependent effect in WT neurons was much smaller compared to Cdr1as-KO, and completely disappeared at DIV21. Network oscillation differences in Cdr1as-KO, which were efficiently modulated by the sustained high expression of miR-7, is a strong indication of an imbalance between glutamatergic and inhibitory signalling in Cdr1as-KO neuronal network.

Finally, to study timepoints where the network reaches its maximal firing peak (maturation of the network) we analyzed network burst peaks by measuring the maximum spike frequency in the average network burst. We did not find significant differences between WT and Cdr1as-KO neurons over time, and we observed a significant down-regulation on the burst peak after miR-7 overexpression affecting equally both genotypes (**Fig. 4C**).

These results suggest that the modulation of miR-7 expression has a great influence on neuronal network activity regulation, but the extent and strength of this influence is modulated by the expression of the Cdr1as. Therefore, in constitutive absence of Cdr1as the system is more prone to the effects of miR-7, especially noticeable in very late times of synaptic maturation.

### Sustained miR-7 overexpression down-regulates Cdr1as and restricts its residual expression to the soma

To better understand molecular mechanisms underlying Cdr1as:miR-7 interactions (and potential interactions with Cyrano), we performed single molecule RNA in situ hybridization (Stellaris, Methods) for Cdr1as (**Fig. 5A**). Controls were Cdr1as-KO cells (**Suppl. Fig.2B**). Additionally, we imaged Cyrano RNA (**Fig. 5C**). Images were quantified (Methods) across biological replicates (independent cultures from different animals). Perhaps surprisingly, we noticed that upon sustained miR-7 overexpression in WT neurons, the distribution of Cdr1as molecules was restricted to neuronal somas (and close proximal space), while massively cleared from all neurites (**Fig. 5A-B**). Additionally, we also observed a reduction of Cyrano molecules. However, this reduction was only significant in the soma but not in neurites (**Fig. 5D)**.

**Figure 5.**
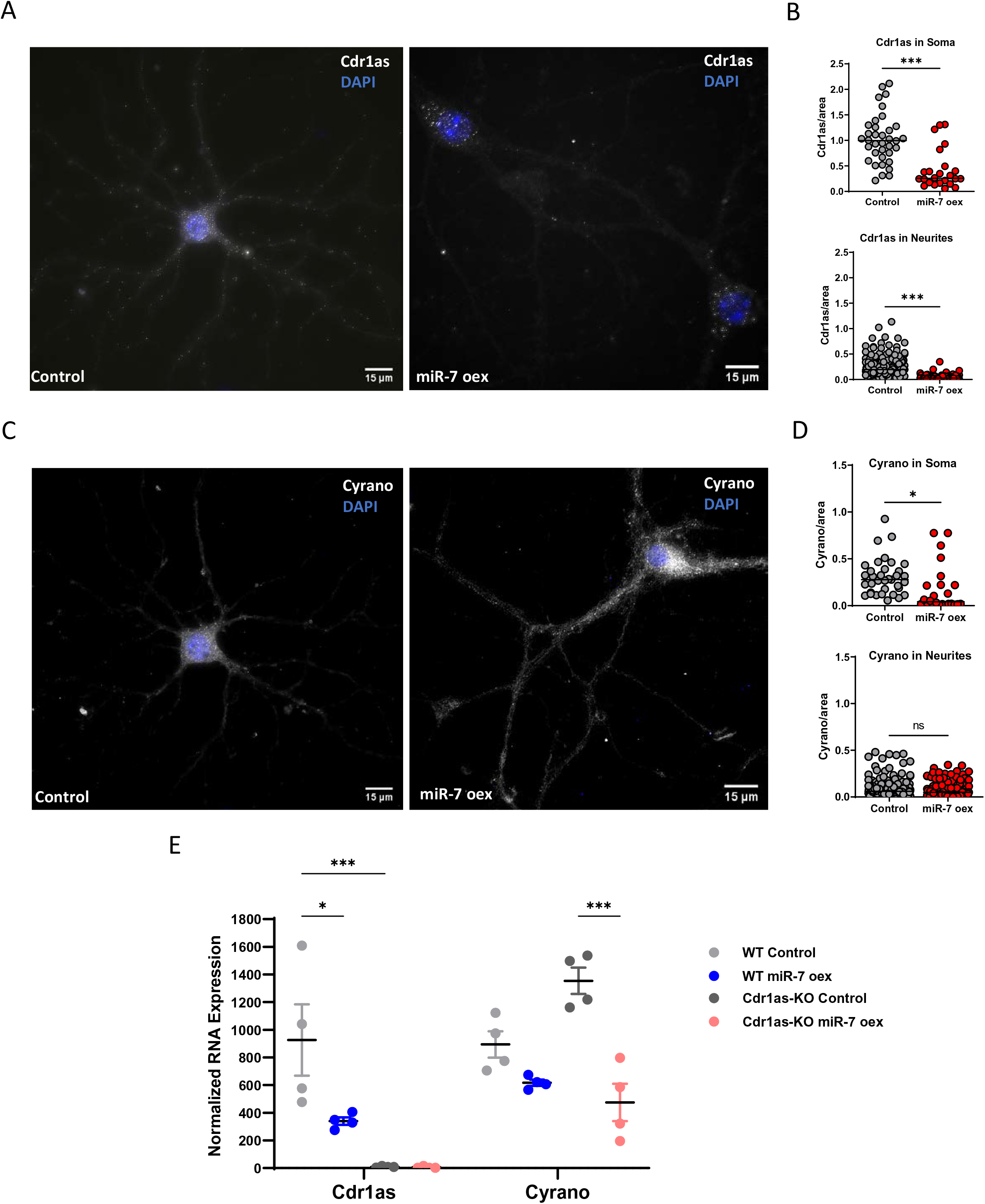
miR-7 overexpression downregulates Cdr1as, removes it from neurites and restricts its residual expression to the soma. **(A)** Single molecule RNA FISH (Stellaris, Methods) of Cdr1as (white) performed in WT neurons DIV21 from neurons infected with control or miR-7a overexpression construct. DAPI: blue. **(B)** smRNA FISH quantification of Cdr1as molecules (following Raj et al., 2008, Methods) ; Each dot represents the mean number of Cdr1as molecules in an independent cell compartment (soma or neurite) normalized by area (100 px = 21,5 um). Control neurons: 25 somas and 110 neurites (2 independent cultures from 2 animals). miR-7 overexpression neurons: 35 soma and 230 neurites (2 independent cultures from 2 animals). Horizontal line: Median. P value: unpaired t-test with Welch correction **(C)** Single molecule RNA FISH (Stellaris, Methods) of lncRNA Cyrano (white) performed in WT neurons DIV21 from neurons infected with control or miR-7a overexpression construct. DAPI: blue. **(D)** smRNA FISH quantification of Cyrano molecules (following Raj et al., 2008, Methods). Each dot represents the mean number of Cyrano molecules in an independent cell compartment (soma or neurite) normalized by area (100 px = 21,5 um). Control neurons: 25 somas and 110 neurites (2 independent cultures from 2 animals); miR-7 overexpression neurons: 35 soma and 230 neurites (2 independent cultures from 2 animals). Horizontal line: Median. P value: unpaired t-test with Welch correction. **(E)** Quantification of Cdr1as and Cyrano (Nanostring nCounter, Methods),performed in WT and Cdr1as-KO neurons DIV21 from neurons infected with control or miR-7a overexpression construct. RNA counts are normalized to housekeeping genes (*Actb, Tubb5 and Vinculin*). Each dot represents an independent biological replicate (4 independent primary cultures from 4 animals). P value: 2-way ANOVA. Error: SD. Horizontal bar: Median. P value: two-way ANOVA, all other comparisons were not statistically significant (not shown).

To investigate if Cdr1as and Cyrano are spatially co-localized in control neuronal conditions or after miR-7 overexpression we used a computational pipeline (Eliscovich et al., 2017) for quantification of single-molecule spatial correlations (Methods). We measured all mutual nearest neighbour distances (MNN) of molecule associations: Cdr1as-Cyrano in comparison with positive and negative controls (Methods). The results from the cumulative distribution of MNN distances between each pair of probes showed no significant association in space between Cdr1as and Cyrano molecules neither in control neurons nor after miR-7 overexpression, as compared to positive molecule-association controls (**Suppl. Fig. 5A**). The association in space was tested in whole cells, neurites, and somas, separately (**Suppl. Fig. 5B**), We did not find significant differences between Cdr1as-Cyrano distances and the negative control in any of the tested compartments, for either of the two tested experimental conditions. Therefore, we conclude that Cdr1as and Cyrano are not specifically co-localized in a biologically relevant distance.

Independent quantification of RNA expression by the Nanostring system, (Methods) confirmed statistically significant decrease of Cdr1as and Cyrano molecules in WT neurons after miR-7 overexpression. We observed that this is also the case for Cyrano in Cdr1as-KO neurons (**Fig. 5E**), after sustained miR-7 overexpression. Altogether, these data indicate that we have identified a new layer of regulation of the Cdr1as-Cyrano-miR-7 axis, where not only expression change of both lncRNAs affect miR-7 levels (Kleaveland et al., 2018; Piwecka et al., 2017), but also exposure to sustained high concentrations of miR-7 generates a negative feedback loop that down-regulates both lncRNAs, and for Cdr1as this decrease happens mainly in neuronal projections. The molecular mechanisms responsible for this down-regulation and whether it is a direct or indirect effect of miR-7 up-regulation remains unknown.

The observations regarding Cdr1as cleared out from neurites in WT neurons, may explain in part the reduced phenotype observed in local and network neuronal activity, and glutamate secretion of WT neurons after sustained miR-7 overexpression. A gradual reduction of miR-7 main regulator from the intracellular compartments closer to synaptic zones, during the time course of miR-7 overexpression, could be associated with the arising of a similar synaptic phenotype as to Cdr1as-KO neurons, but lower in strength depending on the time after miR-7 overexpression, a somewhat an indirect acute knockdown of miR-7 regulators, spatially restricted to neurites.

### Transcriptomic changes caused by sustained miR-7 overexpression are enhanced by the absence of Cdr1as

To further investigate molecular consequences of the interaction of Cdr1as and miR-7, we performed bulk mRNA sequencing of four independent primary neuronal cultures at DIV21 from WT and Cdr1as-KO animals, infected with control or miR-7 overexpression (14dpi). Our identification of differentially expressed genes controlled for batch effects and individual effects (Methods, **Suppl. Fig. 5C**).

First, we investigated predicted miR-7 target genes (Methods). We did not find statistically significant global expression changes of computationally predicted miR-7 target genes when comparing WT and Cdr1as-KO control neurons (**Fig. 6A left panel; Suppl. Table 3**). However, in both, WT and Cdr1as-KO neurons with sustained miR-7 up-regulation, we observed a significant global decrease of miR-7 targets when compared to non-target RNA transcripts. Even more, we noticed that in the case of Cdr1as-KO neurons, miR-7 predicted targets were clearly more strongly down-regulated as compared to WT neurons in the same conditions (significant difference between the shift of cumulative fractions: p value < 1e10^−6^; **Fig. 6A, middle and right panel; Suppl. Fig. 5D, purple and blue lines; Suppl. Table 3**).

**Figure 6.**
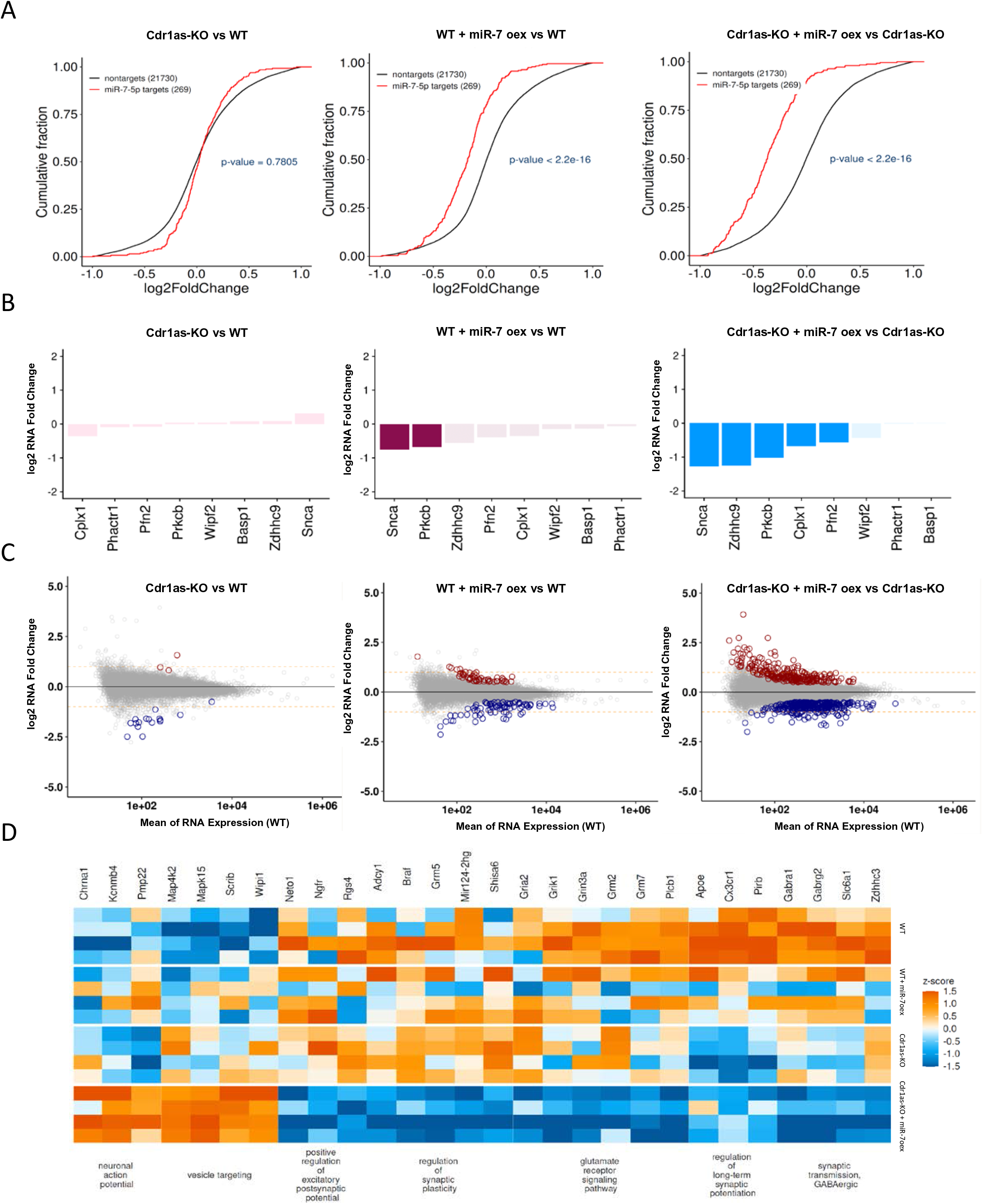
Transcriptomic changes caused by sustained miR-7 overexpression (oex) are enhanced by the loss of Cdr1as and reveal gene regulatory pathways controlled by their interaction. **(A)** Cumulative distribution function (CDF) plot of gene expression comparing all computationally predicted miR-7 targets (Methods, red) to mRNAs lacking predicted miR-7 target sites (non-targets, black), across 4 independent biological replicates of WT, Cdr1as-KO, WT + miR-7 overexpression, Cdr1as-KO + miR-7 overexpression. p value: U Mann–Whitney test. **(B)** miR-7 target genes involved in Insulin granules secretion, experimentally validated by Latreille et al., 2014 in pancreatic β-cells. Bar plots, the mean log2 fold change of 4 independent biological replicates per condition, for each comparison tested. Transparent bars, not significant changes (FDR > 0.05). **(C)** mRNA expression changes for each comparison tested (Cdr1as-KO vs WT, WTmiR-7oex. vs WT, Cdr1as-KO miR-7oex.vs Cdr1as-KO). Plotted is the mean change of 4 independent biological replicates per condition. Red dots: statistically significant up-regulated genes. Blue dots: statistically significant down-regulated genes (Methods). **(D)** Heatmap of statistically significant DE genes associated with GO terms significantly enriched specifically in Cdr1as-KO + miR-7 overexpression vs Cdr1as-KO comparison (specifically up- and down-regulated). Each row represents one independent biological replicate. Z-scores across samples were calculated based on normalized and batch-corrected expression values.

Because miR-7 sustained up-regulation was equally efficient in both genotypes (**Fig. 2E and Suppl. Fig. 3D**), our data suggest that the interaction between Cdr1as and miR-7 is responsible for the differences in observed miR-7 targets regulation strength. This indicates that the constitutive loss of Cdr1as is sensitizing the neurons to the action of miR-7 on its mRNA targets. Predicted targets from a control miRNA were not altered in any of the tested comparisons (i.e., miR-122-5p, miRNA not expressed brain; **Suppl. Fig. 5E**).

Several predicted miR-7 target genes have been experimentally validated as direct miR-7 targets that operate as regulators of secretion in pancreatic cells and recently in hypothalamus (both of them thoroughly studied endocrine cell tissues and highly levels of miR-7 expression; (Bravo-Egana et al., 2008; Kredo-Russo et al., 2012; Landgraf et al., 2007; LaPierre et al., 2022; LaPierre & Stoffel, 2017). Out of these validated miR-7 target genes, *Snca*, a gene encoding for a pre-synaptic scaffold protein and involved in many neurodegenerative diseases (Bennett, 2005), was the most down-regulated miR-7 target gene after miR-7 overexpression regardless of Cdr1as expression, while not being statically significant in controls (**Fig. 6B**).

Strikingly, in numerous cases we observed that experimentally validated secretion-related miR-7 targets that function in pancreatic insulin secretion (Latreille et al., 2014) are exclusively and significantly down-regulated in Cdr1as-KO neurons after sustained miR-7 overexpression (**Fig. 6B, right panel**). Examples include: central regulator of vesicle fusion and SNARE activity (*Cplx1*), cytoskeleton remodeler (*Pfn2*) and an enzyme responsible of regulation of dendritic growth and inhibitory synapses formation (*Zdhhc9*) (**Fig. 6B, right panel**). In contrast, in Cdr1as-KO control neurons we observed only few mRNAs with significantly changed expression compared to WT neurons, (**Fig. 6C; left panel**; **Suppl. Table 2**). Together these observations suggest that the role of miR-7 as a regulator of secretion is conserved between secretory glands (i.e insulin secretion) and forebrain neurons (glutamate secretion), with the crucial difference that the strength of miR-7 action in forebrain neurons depends on Cdr1as levels.

Besides the strong dependence of miR-7 target genes expression on Cdr1as expression, our transcriptomic data revealed many other genes which are dysregulated upon Cdr1as and/or miR-7 perturbation (**Fig. 6C**). Of course, indirect effects are to be expected, but the vast increase in dysregulated genes when combining miR-7 overexpression in Cdr1as-KO neurons points at a synergetic regulation exerted by both RNAs (**Fig. 6C, right panel; Table 2**).

For example, 24.4% of all up-regulated genes (132) are exclusively changing on Cdr1as-KO neurons after miR-7 overexpression (**Fig. 6C, right panel and Suppl. Fig. 6A: upper diagram, grey bubble**). This group of genes does not belong to predicted miR-7 target mRNAs and their gene ontology enrichment analysis linked them with vesicle trafficking and membrane related proteins, and with membrane receptor pathways (**Suppl. Fig. 6B-C-D**), Similarly, 23.3% of all down-regulated genes (151) are only regulated in Cdr1as-KO and their most overrepresented ontology term associate them with plasma membrane proteins with catalytic activity (**Suppl. Fig. 6B-C-D**). On the contrary, only a small number of genes are regulated by miR-7 specifically in WT neurons (3.9% up-regulated −0.6% down-regulated; **Suppl. Fig. 6A, yellow bubbles**).

In summary, our data suggests that global transcriptomic changes caused by miR-7 overexpression are enhanced by the constitutive loss of Cdr1as, connecting RNA changes in neurons with changes in local and network neuronal activity and glutamate secretion. We next turned into a more detailed analysis of specific molecular pathways that might explain these differences.

### Identification of gene regulatory pathways controlled by the interaction of miR-7 and Cdr1as

We performed gene ontology (GO) enrichment analysis with all differentially expressed genes on our transcriptomic data. We identified statistically significant enriched or depleted GO terms in each of the 4 conditions tested (WT, Cdr1as-KO, WT + miR-7 oex and Cdr1as-KO + miR-7 oex; **Fig. 6D, Suppl. Fig. 6B-C-D and Suppl. Table 1**). Analysis the GO terms of genes exclusively enriched or depleted after sustained miR-7 overexpression in Cdr1as-KO (**Suppl. Fig. 6A, upper and lower diagram, red bubble**: up-regulated= 44.4% and down-regulated= 35.2% of differentially expressed genes) found: (1) the largest and specific enriched GO terms are related to the regulation of neuronal action potentials and gene pathways of synaptic vesicle targeting to destination membranes; (2) the depleted terms include, regulation of excitatory postsynaptic potentials, regulation of plasticity and long-term potentiation (LTP), together with regulation of GABAergic transmission and glutamatergic receptors signaling pathways (**Fig. 6D; Suppl. Fig. 6D; Suppl. Table 1**).

Strikingly, all those GO terms enriched or depleted in WT neurons were, conversely depleted or enriched in Cdr1as-KO neurons with miR-7 overexpression, while WT with miR-7 overexpression and Cdr1as-KO control represent intermediates between these 2 scenarios (**Fig. 6D**), which is similar to what we observed for these same conditions on neuronal activity (**Fig. 3 and 4**). These results suggest that the synaptic phenotypes we observed in Cdr1as-KO neurons after sustained miR-7 up-regulation are, in part, explained by the interaction between miR-7 and Cdr1as, which modulates functional cellular pathways reflected in global transcriptomic changes, and that this interaction is essential to trigger all adequate cellular pathways.

## Discussion

miR-7 predominantly high abundance and secretory role in secretory glands, like the pancreas, (LaPierre & Stoffel, 2017; Latreille et al., 2014), together with the evolutionary close relation between neuronal and pancreatic differentiation lineages (Zhao et al., 2007) and the fact that miR-7 evolved within the bilaterian neurosecretory brain (Christodoulou et al., 2010). Led us to speculate that in mammalian forebrain neurons, miR-7 is also involved in regulating secretion. However, as our data show, in primary forebrain neurons, miR-7 is expressed at low levels (~40 miR-7 molecules per neuron in resting state; **Suppl. Fig. 3A**), while Cdr1as is very highly expressed (~250 molecules/neuron, **Suppl. Fig. 2A**).Intriguingly, this is the exact converse in murine pancreatic cells where Cdr1as expression is extremely low while miR-7 is expressed highly (Xu et al., 2015).

How can one explain these expression patterns? Secretion of synaptic vesicles (neurotransmitters) in neurons has to be fast and adaptive to stimuli. We thus speculated that Cdr1as has evolved in the mammalian brain to function as a cellular surveillance system to control, depending on external stimuli, the action of miR-7 on its mRNA targets involved in regulating secretion.

Indeed, in this study, we demonstrate that neuronal adaptation to strong stimulation of murine forebrain neurons causes a rapid transcriptional up-regulation of the circular RNA Cdr1as, comparable to canonical immediate early genes under the same conditions (**Fig. 1 and Suppl. Fig. 1**), and a post-transcriptional stabilization of miR-7 (**Fig. 1C-D)**.

Moreover, our data show that presynaptic terminals of Cdr1as-KO forebrain neurons have modestly increased release of glutamate in resting conditions but strongly increased glutamate release after strong stimulation trains. This increased secretion of excitatory neurotransmitter is completely reverted by sustained miR-7 up-regulation. (**Fig. 2B-C-F**)

We also show that these changes in glutamate release have functional consequences on real-time synaptic activity properties, both at local and at network levels (**Fig. 3A**). Forebrain neurons with a constitutive loss of the Cdr1as locus, and a mild but statistically significant down-regulation of miR-7 (**Suppl. Fig. 2E**), show up-regulated APs frequencies (**Fig. 3B**), which we could directly associate with the increased release of glutamate from the presynaptic terminals (**Fig. 2**). This accelerated AP activity then allows longer and more frequent neuronal bursting, increasing over time of maturation (**Fig. 3C-D**), which more strikingly reflected in a globally uncoordinated neuronal network connectivity (**Fig. 4**), and potentially alter long-term changes in plasticity responses. All synaptic phenotypes induced by the lack of Cdr1as are reverted by the sustained overexpression of miR-7, similarly to what we observed for glutamate release.

Sustained miR-7 overexpression up-regulated equally (±10 folds) the number of miR-7 molecules in WT and Cdr1-KO neurons, which was sufficient to recover any synaptic dysregulation measured in Cdr1as-KO neurons, while synaptic activity of WT neurons was only mildly or not affected, depending on the plasticity parameter tested (**Fig. 3 and 4**). Furthermore, we observed that this is also true for miR-7-dependent target gene modulations and global transcriptomic changes, which heavily depend on the expression levels of Cdr1as (**Fig. 6A-B-C-D**), suggesting that high expression of Cdr1as is critical to control miR-7-dependent effects on bursting regularity, synchrony and oscillation and therefore the balance of excitatory and inhibitory signals.

In addition, we observed clearance of Cdr1as from neuronal projections (**Fig. 5A**). This could explain the differences in strength of glutamate and neuronal activity phenotypes between Cdr1as-KO and WT neurons after sustained miR-7, and might indicate that the role of Cdr1as as a miR-7 buffer is more relevant in neuronal projections, where a restricted but permanent and fast response to sensory stimuli is needed. Therefore, if neurons evolved a molecular mechanism of surveillance that can rapidly up- or down-regulate mRNAs locally in synaptic vicinities, this mechanism would shape the local modifications on the corresponding synaptic terminals. Some evidence to back up this hypothesis has been presented in a preprint where local dendrite down-regulation of Cdr1as, induces an up-regulation of miR-7 and impaired fear extinction memory (Zajaczkowski et al., 2022).

We propose that one of the potential cellular mechanisms through which miR-7, controlled by Cdr1as, directly regulates long-term changes in neuronal connectivity, can be observed in our measurements of network oscillation (**Fig. 4B**).

In general, oscillation differences account for alternating periods of high and low activity of neuronal communication and changes in network periodicity, which is a hallmark of functional neuronal networks with mixed cell populations of excitatory and inhibitory neurons, such as in forebrain, composed of glutamatergic and GABAergic cells. Network oscillation augmentation in Cdr1as-KO, which are then rescued by the sustained up-regulation of miR-7, are an indication of imbalance between glutamate and GABA signaling affecting Cdr1as-KO forebrain network activity. In fact, this is corroborated by the glutamate secretion assay, where we see the increase excitatory signaling in Cdr1as-KO being rescued by miR-7 expression itself (**Fig. 2F**). Based on the enhanced glutamate presynaptic release observed in Cdr1as-KO (**Fig. 2B-C**), we expected that miR-7 overexpression and the complete absence of Cdr1as should thus have opposite effects on secretion genes. This is the case for *Snca*, which is one of the strongest regulated miR-7 target genes.

In forebrain neurons, activity-dependent presynaptic glutamate release depends positively on the concentration of α-synuclein. This could result from the inhibition of glutamate re-uptake by transporters localized on presynaptic, postsynaptic, and astrocytic endings and/or physical disruption of synaptic vesicles on the presynaptic membrane (Sarafian et al., 2017). More refined computational analyses of our expression data identified other genes that might be interesting to investigate (Suppl. Discussion). Furthermore, we proposed potential key gene regulatory pathways connecting these gene ontology terms with our functional phenotypes (Suppl. Discussion; **Suppl. Fig. 7)**.

The exact involvement of other network partners such as lncRNA Cyrano remains to be studied. We did not observe a statistically significant change in Cyrano after sustained depolarization (**Suppl. Fig. 1A**), which indicates that during neuronal activation the regulation of miR-7 levels are not predominantly regulated by Cyrano. We speculate that Cdr1as is the mir-7 regulator dominating miR-7 cellular availability, due to its prevailing stabilization of miR-7. Other less explored players, like Hu RNA binding proteins, which regulate miR-7 processing in non-neural cells through direct binding to the pri-miR-7 could also be involved (Lebedeva et al., 2011). On the other hand, we observed a general down-regulation of Cyrano after sustained overexpression of miR-7 (**Fig. 5E**), but this is happening only in somas, with not much alteration of Cyrano molecules in neuronal projections (**Fig. 5C-D**). Therefore, we confirmed the direct regulation of miR-7 over Cyrano, as suggested before by Kleaveland et al. (2018). We show that miR-7 itself is capable of regulating Cyrano expression, only significantly in soma. To understand better the dynamics of miR-7 regulation it will be necessary to test perturbations on Cdr1as and Cyrano during shorter and longer stimulations.

We summarize our data to propose a molecular model (**Fig. 7**) for the mechanism by which Cdr1as and miR-7 exert their function. In resting neurons (**Fig. 7A**) the removal of Cdr1as causes mild but significant reduction of mir-7 levels and minor effects on glutamate pre-synaptic release clearly reflected in neuronal network modulations. However, the low expression of miR-7 relative to the large amount of Cdr1as impede the continuous action of mir-7 on secretion targets.

**Figure 7.**
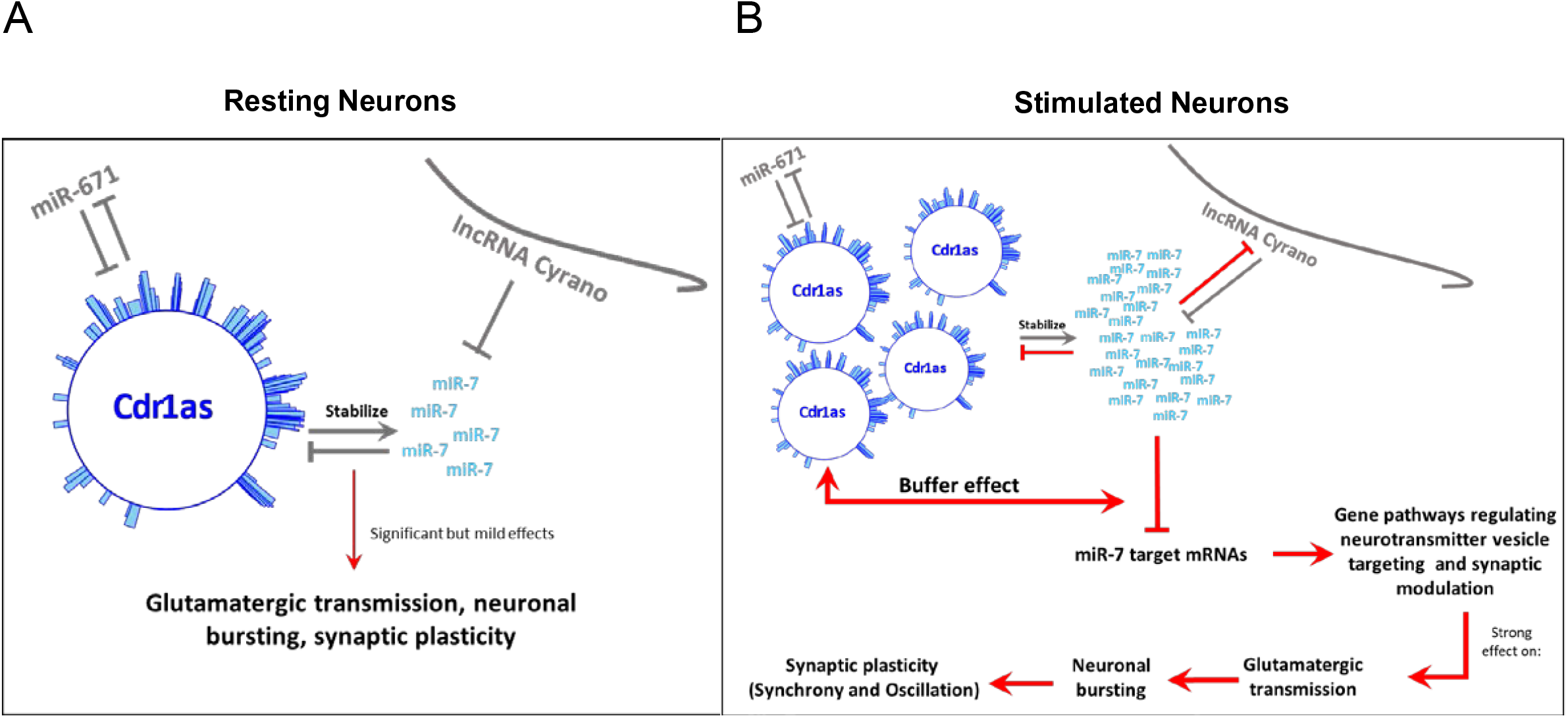
Proposed coordinated mechanism and function of Cdr1as and miR-7 in resting and stimulated neurons. Grey arrows: Published data (Hansen et al., 2011; Memczak et al., 2013; Hansen et al., 2013; Piwecka at al., 2017; Kleaveland et al., 2018). Red Arrows: New interactions propose in this study. **(A)** Binding sites of miR7 to Cdr1as (blue bars) are drawn following Piwecka et al. 2017. In resting neurons upon removal of Cdr1as, mir-7 levels are reduced and there are mild but significant effects on glutamate pre-synaptic release, which induce changes in synchrony and oscillation of the neuronal network. However, the low expression of miR-7 relative to the large amount of Cdr1as impede the continuous action of mir-7 on its targets. **(B)** Strong sustained neuronal stimulation transcriptionally up-regulates Cdr1as which prevents miR-7 degradation and therefore rapidly post-transcriptionally stabilizes miR-7. This persistent increase in miR-7 will cause long-lasting down-regulation of Cdr1as from neurites, so that the buffer can self-regulate and therefore dynamically modulate glutamatergic transmission and neuronal network activity. We propose that the post-transcriptional nature of this buffer system allows fast and local regulation of vesicle targeting in synapses.

During strong sustained neuronal stimulation (**Fig. 7B**) Cdr1as is transcriptionally up-regulated which prevents miR-7 degradation and therefore, rapidly post-transcriptionally stabilizes miR-7. Nevertheless, miR-7 long-lasting persistent increase will down-regulate Cdr1as from neurites, so that the buffer can self-regulate and therefore dynamically modulate glutamatergic transmission and neuronal network activity. Overall, we hypothesize that the post-transcriptional nature of this buffer system allows fast and local regulation of vesicle targeting in synapses and ultimately synaptic plasticity.

## Acknowledgments

We thank all members of the Rajewsky lab, past and present, for helpful discussions and advice. We thank Margareta Herzog for all organizational help. CACJ thanks Christine Kocks and Anastasiya Boltengagen for technical advice on single molecule FISH, Thomas Müller for help with primary neuronal cultures, Ivano Legnini for advice on neuronal stimulation experiments, Ilan Theurillat for feedback on manuscript edition and Petra Stallerow for technical support with animal caretaking. SJK thanks Marcel Schilling and Marvin Jens for computational analysis assistance. EG thanks Marie Piraud. NR thanks DFG Leibniz Award and DFG Neurocure/BrianBank for support. CACJ was funded by the MDC graduate program, DZHK project 81X2100155 and DFG Neurocure/BrianBank. SJK was funded by EU ITN – circular RNA Biology Training Network: circRTrain (721890) and DFG Excellence Cluster (DFG EXC2049). GZ was funded by DFG Leibniz Award. EG was funded by Helmholtz Association’s Initiative and Networking Fund through Helmholtz AI. MP has been supported by the Polish National Agency for Academic Exchange (Polish Returns grant no. PPN/PPO/2019/1/00035/U/0001) and the National Science Centre (grant no. 2018/30/E/NZ3/00624).

## Author contributions

CACJ and NR conceived the study. CACJ performed all the experiments. SJK performed all bioinformatic analysis with exception of computational RNA colocalization performed by GZ. GT supported CACJ with sequencing libraries, animal handling and animal maintenance. CACJ, ZF and AW performed glutamate imaging experiments. ZF and AW performed all analysis of glutamate imaging. SJK and EG performed random forest analysis. MP and NR supervised CACJ in the beginning of the project. NR supervised CACJ throughout whole study. CACJ and NR wrote the manuscript with inputs from all authors.

## Methods

### I. Animals

#### Cdr1as knockout mice

Cdr1as knockout (Cdr1as-KO) strain was generated and maintained on the pure C57BL/6N background. All animals used in this work came from intercrosses of hemizygous Cdr1as-/Y males and heterozygous Cdr1as+/− females in (F3 generation from original founder) and mutants were compared to wildtype littermate controls. Animals were kept in a pathogen-free facility in a 12 h light–dark cycle with ad libitum food and water. Animal care and mouse work were conducted according to the guidelines of the Institutional Animal Care and Use Committee of the Max Delbrück Center for Molecular Medicine, the Landesamt für Gesundheit und Soziales of the federal state of Berlin (Berlin, Germany).

### II. Biochemical and Molecular Biology

#### Cloning, transformation, and propagation of plasmids

Cloning was performed according to standard methods. Plasmids were cut using appropriate restriction enzymes (Fast Digest-Thermo Scientific), dephosphorylated with FastAP and purified from an agarose gel with a Zymoclean Gel DNA Recovery kit (Zymo Research). Inserts were either PCR amplified, then digested with the same set of restriction enzymes or oligonucleotides with fitting overhangs were annealed and phosphorylated with T4 PNK. Insert and cut plasmid were ligated with T4 DNA Ligase and subsequently transformed into chemically competent bacteria (Mix&Go Competent Cells, DH5 α) in presence of the corresponding antibiotic for selection. Finally, plasmids were isolated using ZymoPURE Plasmid Miniprep or ZymoPURE Plasmid Midiprep kit from bacterial culture. For plasmids obtain from Addgene, same procedure of propagation and plasmid purification was used.

#### Genotyping of mutant animals

To genotype Cdr1as KO strain, genomic DNA was extracted from tail cuts taken from newborn or adult animals. Tissue was digested using QuickExtractTM DNA Extraction Solution 1.0 at 65°C for up to 30 mins depending on the tissue size, reaction was stopped at 98°C for 2 mins and 1ul of the supernatant, containing the genomic DNA, was used for the follow up end-point PCR detection of wildtype or mutant splice sites splice sites. Genotypes were determined by end-point PCR performed on corresponding template genomic DNA using a set of primers for 3’SS multiplex detection (3’SS_For + 3’SS_Rev + 5’SS+For). PCR products for Wildtype (355bp), Heterozygotes (418 and 355 bp) and Knock-out (418bp) animals were analyzed on a 2.5% agarose gel.

#### Nanostring

To achieve direct quantification of targeted single RNA molecules in a sample of total RNA. 100ng of RNA were incubated with a 72-plex Core Tag set, diluted custom-made probe mix (reporter + capture probes) and corresponding hybridization buffer, at 67°C for 18h, followed by a cool down step at 4°C for 10mins. All samples were diluted to a final volume of 30 ul and loaded in a nCounter gene expression cartridge (12 sample panel). Quantification of RNA molecules was done by nCounter SPRINT^TM^ Profiler (NanostringTechnologies, USA) instrument according with manufacturer run instructions. Normalized counts of RNA molecules were obtained using nSolver^TM^ Data analysis software.

#### Overexpression miR-7

The sequence of pri-miR-7a was amplified from mouse brain tissue and cloned into an AAV vector (Addgene Plasmid #99126), driven by a neuronal promoter (hSyn1) and expressed together with mCherry fluorescent reporter. The final viral particles (AAV serotype 9 and mutated versions of Serotype 9) were generated by Charité Viral Core Facility. An AAV that only expressed the mCherry protein under the same neuronal promoter was use as control condition (Addgene Plasmid #114472). Wildtype and Cdr1as-KO neurons were infected at DIV4-6 by directly adding the viral particles into the culturing media at a titer of 10^9^ VG/ml. half of the media was exchange after 5 days of infection and then once a week. At DIV21, cells were fixed or harvested in Trizol according to the different downstream experiments.

#### Quantitative real time PCR (qPCR)

Quantitative real time PCR experiments were conducted on an ABI StepOne Plus instrument or on a Roche LightCycler 96 System. cDNA was diluted up to 20X with ddH2O and mixed with 2x Biozym Blue S’Green + ROX qPCR Master Mix and 1μl of 10μM Primer mix to obtain 20μl of total volume per well. All measurements were conducted at least in 3 technical replicates. For miRNA expression analysis, cDNA was diluted 3X with ddH2O and TaqMan assays (Applied Biosystems) with an attached FAM dye were quantified using TaqMan Universal master Mix II, no UNG-1 (Applied Biosystems). Relative quantification of gene expression was performed by using the comparative ΔΔCT method (Pfaffl, 2001).

#### Reverse Transcription

Total RNA was reverse transcribed to cDNA using Maxima H Minus Reverse Transcriptase (Thermo Scientific) (100 U/μg RNA). First, 0.5 −1 μg of total RNA was mixed with 1 mM dNTPs, 0.5 μg/μl random hexamers, denatured at 65°C for 5 min and then placed on ice. Immediately, 200 U/μl of Reverse Transcriptase, 4 μl 5x RT Buffer and 0.5 μl Ribolock (40U/μl) were added. The cDNA synthesis was carried out as follows: 10 min at 25°C, 1h at 50°C, 5 min at 85°C. For miRNAs, instead of normal random hexamers, gene-specific stem loop primers (Taqman RT primers) were used, which only need a short overlap with the miRNAs 3’end. First, 100 ng of total RNA was reverse transcribed using SuperScript III (Invitrogen) (100 U/reaction), 1 mM dNTPs, 20 U Ribolock (40U/μl), 1x TaqMan RT primer per each gene of interest (miR-7a, miR-671, Let-7a, snoRNA202, U6 snRNA) (Applied Biosystems) and 1x first-strand synthesis buffer were filled up to 20 μl reaction volume with ddH2O and mixed. Then the gene-specific cDNA synthesis was carried out as follows: 30 min at 16°C, 30 min at 42°C and terminated for 5 min at 85°C.

#### RNA extraction

Total RNA from primary neurons was extracted using home-made TRIzol reagent in combination with Direct-zol RNA Kit (Zymo Research). The samples were fixed in cold methanol for 10 mins at −20C, then washed in cold PBS, TRIzol was directly added to each well and cells were scraped away to ensure full detachment of all neuronal processes. The dissociated sample was collected in a tube with silica beads and homogenized at 5500 rpm 2x for 20secs. Then equal volume of 100% Ethanol was added directly to the sample and then transferred to a Zymo-Spin^TM^ IIICG Column and centrifuged (30secs x 16000g). The flowthrough was discarded and DNAse I (Zymo Research) treatment followed, incubating the column with 6U/ul for 15 mins at room temperature. Afterwards, 3 consecutive washing steps were performed and finally the RNA was eluted from the Column in DNAse/RNAse-free water.

### III. Imaging techniques and analysis

#### smRNA FISH (Stellaris) (Raj et al., 2008)

Single molecule fluorescent *in situ* hybridization (smFISH) protocol was performed in Wildtype and Cdr1as-KO primary neurons after 14-21 DIV. Stellaris oligonucleotide probes complementary to Cdr1as, Cyrano (Oip5-as1), Hprt, circHipk3 or linHipk3 were designed using the Stellaris Probe Designer (LGC Biosearch Technologies) as conjugates coupled to Quasar 670 or Quasar 570. Neurons were fixed in 4% Formaldehyde for 10 mins. and probes hybridized overnight in 10% formamide at 37°C and 100 nM, washed according to Stellaris protocol (Biosearch Technologies), DAPI 5 ng/ml in second wash and mounted with Gloxy buffer. Images were acquired on an inverted Nikon Ti-E microscope with 60x oil NA1.4 objective and Andor iXON Ultra DU-888 camera; Z stacks had 0.3 μm spacing and were merged by maximum intensity projection. Dot detection was performed using StarSearch software from Raj Lab (https://github.com/arjunrajlaboratory/rajlabimagetools/wiki) on manually segmented neurons (somas and neurites). Posterior quantification analysis was done counting the number of dots of Cdr1as or Cyrano molecules normalized by area and per compartment. All statistics were calculated using Mann-Whitney test, nonparametric rank comparisons, between different conditions (GraphPad Prism 9.1.0), statistical significance was assigned for P<0.05

#### Single-molecule miRNA FISH (ViewRNA Cell Plus)

Single molecule miRNA fluorescent *in situ* hybridization (miRNA smFISH) protocol was performed in Wildtype and Cdr1as-KO using primary neurons after 21DIV. miRNA probes for mature sequence of miR-7a-5p were obtain from available commercial catalog (Thermo scientific) and label probes for A670 were used. miRNA smFISH protocol was performed according to manufacturer protocol (ViewRNA Cell Plus – ThermoFisher) plus the addition of protease K (Invitrogen) treatment in a dilution 1/1000 for 10 mins, before miRNA probe hybridization step. Finally, the samples were mounted with Prolong Gold Antifade mounting media (Invitrogen) and imaged using Keyence BZ 9000 or Leica Sp8 confocal microscope. Image processing was done using Fiji-imagej.

#### smRNA FISH colocalization analyses

Cdr1as and lncRNA Cyrano molecules within neurons, we performed colocalization analyses of single molecules based on MNN distances and probability of random association calculation. smRNA FISH (Stellaris) images of Cdr1as and Cyrano were used as the test condition, Cdr1as and Tfrc (transferrin receptor C) as negative control and images of two probes within the same RNA molecule (linHipk3 and circHipk3) as positive control of true colocalization. Dot detection was performed using StarSearch software from Raj Lab (https://github.com/arjunrajlaboratory/rajlabimagetools/wiki) on manually segmented neurons (somas and neurites). The analyses to measure Intermolecular Distances and Determining the Significance of association, were done based on the work of (Eliscovich et al., 2017), but rewrite in a new R script according to our requirements.

#### Synaptic Glutamate Release

##### i) iGluSnFR

AAV1 particles containing a Glutamate sensor under the human synapsin-1 promoter (pAAV.hSyn.iGluSnFr.WPRE.SV40 (Borghuis et al., 2013) were obtained from A.W Lab at MDC Berlin. Primary neuronal cultures were infected at DIV 4-6 with the AAV particles to express the glutamate sensor at the cell surface and then recorded at DIV17-21.

##### ii) Image acquisition

All image acquisition was done as previously described in Farsi et al. (2021) (Farsi et al., 2021) using same instruments, solutions and recording protocols. In brief, neurons were incubated at room temperature in Tyrode’s buffer (120 mM NaCl, 2.5mM KCl, 10 mM glucose, 10 mM HEPES, pH 7.4, osmolality was adjusted to that of the culture medium with sucrose). Action potential-evoked glutamate release was recorded in a chamber with two electrodes for electrical stimulation, mounted on a Nikon Eclipse Ti inverted microscope equipped with a PFS focus controller, a prime 95B scientific CMOS (Photometrics) camera and a pulse generator (HSE-HA, Harvard Apparatus). Emission was collected with a 60x, 1.49 NA Nikon objective. Evoked glutamate release was performed by continuous imaging at 20 Hz after application of20 field stimuli at 0.5 Hz in the presence of 50 μM AP5 (l-2-amino-5-phosphonovaleric acid) and 10 μM CNQX, to block postsynaptic, imaging solution was supplemented with 2 mM CaCl2 and 2 mM MgCl2. Spontaneous events were then captured by 5-min continuous imagingat 20 Hz at 4mM CaCl2, in the presence of 0.5 μM TTX to block action potential firing.

##### iii) Image analysis

All image analysis was done as previously described in Farsi et al. 2021. In brief, time-lapse images were loaded in MATLAB (Mathworks, Natick, MA) and after de-noising and filtering, active synapses were detected as local maxima on the first derivative over time of the image stack. Regions of interest (ROIs) were defined by stretching the maxima to a radius of 4 pixels (700 nm). Taking the first derivative over time allowed to resolve fluorescence changes associated with evoked or spontaneous release and localize release sites.

##### iv) Release probability calculation

As previously described in Farsi et al. 2021, to characterize the release properties of individual synapses, filtered fluorescence traces were used for peak detection. Peaks with the amplitude seven times greater than standard deviation of baseline trace during spontaneous imaging were counted as a successful glutamate release event. Spontaneous frequency was calculated from the total number of events detected during 5-min acquisition. Evoked probability for each synapse was obtained by dividing the number of events happening within one frame after stimulation by the total number of stimulations.

##### v) Kernel Density Estimation

All release probability calculations were plotted using the Kernel-Density-Estimate and creating a curve of the distribution of individual measurements. The curve is calculated by weighing the distance of all the points in each specific location along the distribution. Each data point is replaced with a weighting function to estimate the probability density function. The resulting probability density function is a summation of every kernel.

### IV. Cell Culture Methods

#### Primary Neuronal culture

Primary cortical neurons were prepared from C57BL/6N mouse pups (P0) as previously described in Kaech and Banker, 2007 (Kaech & Banker, 2006). In brief, cortices were isolated from P0 C57BL/6N mice from Wildtype and Cdr1as-KO genotypes. The tissue was dissociated in papain at 37C and then the single cell suspension was seeded in a previously coated glass coverslips, to finally place them on top of a monolayer of feeder astroglia. The cells were maintained in culture up to 21 days in Neurobasal-A medium (Invotrogen) supplemented with 2% B-27(Gibco), 0.1% PenStrep (10 kU/ml Pen; 10 mg/ml Strep) and 0.5 mM GlutaMAX-I (Gibco) at 37°C and 5% CO_2_.

#### K+ treatments

On DIV13 neurons were incubated over night with a mix of: 0.5μM TTX (tetrodotoxin), 100μM AP5 (l-2-amino-5-phosphonovaleric acid) and 10 μM CNQX, to silence all spontaneous neuronal responses. On DIV14 neuronal media was exchange for stimulation media (170mM NaCl, 10mM HEPES pH7.4, 1mM MgCl2, 2mM CaCl2), containing whether (1) 60mM KCl, (2) 50μM DRB (5,6-Dichloro-1-β-D-ribofuranosylbenzimidazole) or just (3) NaCl as osmolarity control. Cells (neuronal cultures and astrocyte-feeder layer) were incubated at 37°C; 5%CO2 for 1 hour, fixed with cold methanol for 10 mins. and used immediately for RNA extraction.

#### Glutamate Secretion assay

To assess the levels of secreted Glutamate in Wildtype and Cdr1as-KO neurons with or without miR-7a overexpression, we implemented the luciferase-based Glutamate-Glo™ assay. Neurons were plated in 48-well at equal confluency and media from each well was collected at DIV18-21 and saved at −20°C until ready to perform the assay. 25μl of media from each sample were transferred into a 96-well plate to perform the final reaction. A negative control of only buffer was included for determining assay background and a control of no-cells mediawas used to determine basal levels of Glutamate. Samples were mixed with GlutamateDetection Reagent prepared according to manufacturer instructions and the plate incubated for60 minutes at room temperature. Recording of luminescence was done using a plate-reading luminometer (Tecan M200 infinite Pro plate reader) following manufacturer protocol. A stocksolution of glutamate 10mM was used to create the standard curve (100μM to 1.28×10^−3^μM) and as positive control. All Glutamate concentrations were estimated based on standard curve linear regression. Analyses were done by comparison of RLU values across conditions and allstatistics were calculated using two-way ANOVA, between different conditions, statistical significance was assigned for P<0.05.

#### Multi-Electrode Array (MEA) Recordings

Cells were seeded in 48-well CytoView MEA plates with 16 poly-3,4-ethylendioxythiophen (PEDOT) electrodes per well (Axion Biosystems). Each well was coated with 100 μg/mL poly-D-Lysine and cells were seeded to a density of 100-150K mixed with 1 μg/mL laminin (FUDR was added to the media one day after seeding). Neurons were grown in supplemented Neurobasal-A media for the first 7 days, then half of the media was changed to BrainPhys^TM^, supplemented with B-27, one day before each recording. Recordings of spontaneous extracellular field potentials in neurons from DIV 7 to DIV21 were performed using a Maestro MEA system and AxIS Software with Spontaneous Neural Configuration (Axion Biosystems, Atlanta, GA, USA). Spikes were detected using an adaptive threshold set to 6 times the standard deviation of the estimated noise. The plate was first allowed to ambient in the Maestrodevice for 3 minutes and then 10 minutes of raw spikes data was acquired for analysis.

##### i) MEA Data analysis

For MEA data analysis we used a sampling frequency of 12.5 kHz and the active electrode selection criteria was 5 spikes/minute. (1) Mean firing rate (MFR): action potentials (APs) frequency per electrode. (2) Burst Frequency: Total number of single-electrode bursts divided by the duration of the analysis, in Hz. (3) Burst Duration: Average time (sec) from the first spike to last spike in a single-electrode burst. (4) Network Asynchrony: Area under the well-wide pooled inter-electrode cross-correlation, according to Halliday et al., 2006. (5) Network Oscillation: Average across network bursts of the Inter Spike Interval Coefficient of Variation (standard deviation/mean of the inter-spike interval) within network bursts. (6) Burst Peak: peak of the Average Network Burst Histogram divided by the histogram bin size to yield spikes per sec (Hz). All statistics were calculated using 2-way ANOVA mixed model with Bonferroni correction for multiple measurements, between different conditions, statistical significance was assigned for P<0.05

### V. Bulk RNA-Sequencing

#### Total RNA libraries and sequencing (circRNA detection)

1 μg of total RNA was used as starting material for total RNA libraries and spiked with from Wildtype and Cdr1as-KO neurons. Then, ribosomal RNA (rRNA) was depleted with the Ribominus Eukaryote Kit v2 or with a RNase H approach (Adiconis et al., 2013). Successful ribodepletion was assessed by Bioanalyzer RNA 6000 Pico Chip. RNA was fragmented and subsequently prepared for sequencing using Illumina Truseq Stranded Total RNA Library Prep kit. Libraries were sequenced on a Nextseq 500, at 1×150 nt.

#### PolyA+ libraries and Sequencing

All samples from Wildtype and Cdr1as-KO neurons, control and miR-7 overexpression, were prepared as described in the TruSeq Stranded RNA sample preparation v2 guide. In brief, 500ng of RNA were fragmented, reverse transcribed and adapter-ligated, followed by a pilot qPCR to determine the optimal PCR amplification. Library quality was assessed using Tapestation (DNA1000 kit) and Qubit. Samples were sequenced on an Illumina NextSeq 500 with 1×150 bp.

#### Small RNA libraries and Sequencing

All samples from WT and Cdr1as-KO neurons were prepared from total RNA as described in Illumina TruSeq Small RNA Sample Prep Kit and sequenced on a NextSeq 500 with 1×50 bp.

#### RNA-Seq analysis

RNA-seq reads were mapped to the mouse mm10 genome assembly using STAR v2.7.0a (Dobin et al., 2013). Aligned reads were assigned to genes using annotations from Ensembl (Mus_musculus, release 96) and featureCounts v1.6.0 () with the parameter reverse stranded mode (−s=2). Differential gene expression analysis was done using DESeq2 v 1.30.1(Love et al., 2014), taking both batch effects and nested effect into account. Specifically, we used multi-factor design that consider the batch effect to WT vs KO and WT vs KOm7oe (design = ~ 0 + group + batch), while used nested effect to WT vs WTm7oe and KO vs KOm7oe in paired experiments (design = ~genotype+genotype:batch+genotype:cond). Significance threshold of differential expression genes was set to adjusted P-value of 0.05 and log2-fold-change of 0.5.

#### miR-7-5p target regulation analysis

To test if log2FoldChange distribution of miR-7 targets are significantly different to other genes used in differential expression genes analysis, we performed a two-sided Mann-Whitney U-test for each comparison. A list of predicted conserved targets of miR-7-5p and miR-122 as a control were downloaded from TargetScan Mouse, release 7.2 (Agarwal et al., 2015) and miRDB (Chen & Wang, 2020). We only used targets exist in both databases.

#### Gene ontology (GO) analysis

Gene ontology enrichment analysis was done using topGOtable function in the pcaExplorer (Marini et al., 2019). In GO analysis, genes showing average log2 fold change of 0.5 and adjusted p value less than 0.05 were considered significant and all expressed genes were used as background. We used the elim algorithm instead of the classic method to be more conservative and excluded broad GO terms with more than 1000 listed reference genes.

#### Gene networks analysis

To investigate whether genes related to neurological phenotypes are regulated by noncoding RNAs (in particular miR-7) and to find a subset of genes whose expression profiles are the most predictive of a target gene’s expression profile, we used randomForest approach (Breiman, 2001). To find direct and indirect targets of miR-7, we could evaluate the learned direct targets from our GRN structure, we do network learning for gene lists provided by prior knowledge and indirect targets of miR-7 and indirect targets of miR-7 and direct targets of miR-7 separately. To narrow down gene list in the GRN, we filtered genes that overlapped between two or more genes of interest were selected as key regulator genes for each run. Leave-one-sample-out cross-validation was used to train random forest models and evaluate their prediction performance. Prediction performance was evaluated by root mean squared error (RMSE), a standard measure in the evaluation of quantitative predictions, which has aminimum of zero and no upper bound. In addition, we computed differences of the predicted or ground truth values relative to the training examples and computed the cosine similarity of these two difference vectors, which is 1 if predictions and ground truth show consistent patternsand −1 if predictions and ground truth show opposite patterns. The prediction performance on left-out test samples was good, which implies that the models learned some meaningful relationships. To validate our prediction, we downloaded protein-protein interaction from stringDB (10090.protein.info.v11.5).

---

**Supplementary Figure 1.**
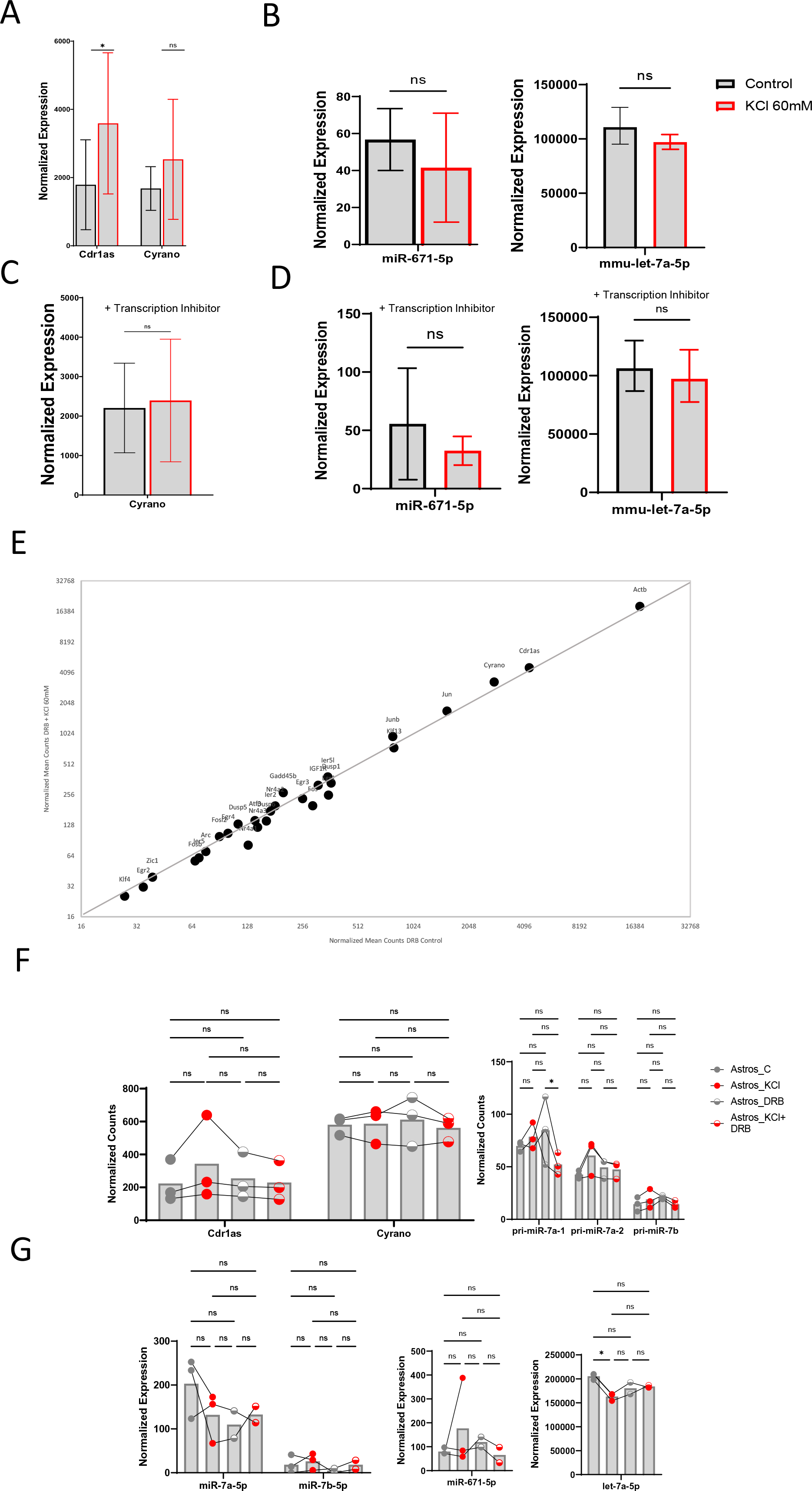
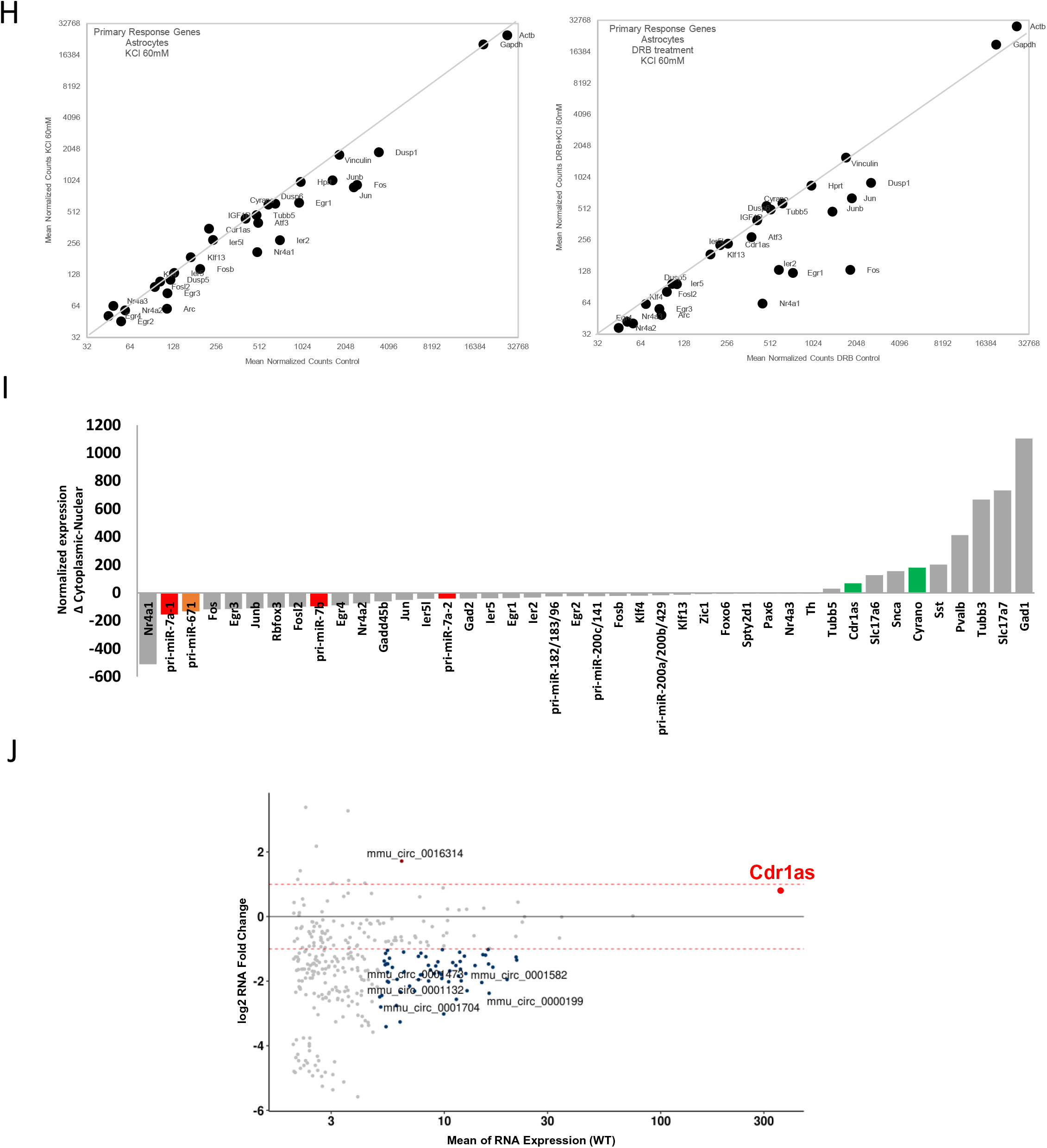
(A) Quantification of Cdr1as and Cyrano (Nanostring nCounter, methods), before and after sustained depolarization. RNA counts are normalized to housekeeping genes (Actb, Tubb5 and Vinculin). Bar plot represents the mean of 4 biological replicates (3 independent primary cultures from 3 animals). P value: ratio paired t-test. Error: SD. (B) Expression levels of mature miR-671 (left) and let-7a (right) quantified by small RNA-seq for 3 independent primary cultures, before and after sustained depolarization. Bar plot represents the mean. P value: ratio paired t-test. Error: SD (C) Quantification of Cyrano (Nanostring nCounter, methods), before and after sustained depolarization plus pre-incubation with transcription inhibitor (DRB). RNA counts are normalized to housekeeping genes (Actb, Tubb5 and Vinculin). Bar plot represents the mean of 3 biological replicates (3 independent primary cultures from 3 animals). P value: ratio paired t-test. Error: SD. (D) Expression levels of mature miR-671 (left) and let-7a (right) quantified by small RNA-seq for 3 independent primary cultures, before and after sustained depolarization plus pre-incubation with transcription inhibitor (DRB). P value: ratio paired t-test. Error: SD (E) RNA Quantification before and after K+ treatment depolarization plus pre-incubation with transcription inhibitor (DRB) (Nanostring nCounter, methods). Cdr1as and Cyrano shown in red. RNA counts are normalized to housekeeping genes (Actb, Tubb5 and Vinculin). Each dot represents the mean of 3 biological replicates (3 independent primary cultures from 3 animals). Grey line: linear regression. (F) Quantification of Cdr1as, Cyrano and primary miRNAs (Nanostring nCounter, methods), in primary astrocytes exposed to neuronal culture (feeder layer) before and after sustained depolarization plus pre-incubation with transcription inhibitor (DRB). RNA counts are normalized to housekeeping genes (Actb, Tubb5 and Vinculin). Bar plot represents the mean of 3 biological replicates (3 independent primary cultures from 3 animals). P value: two-way ANOVA. (G) Expression levels of mature miR-7 isoforms (left) miR-671 (middle) and let-7a (right) quantified by small RNA-seq for 3 independent primary cultures, in primary astrocytes exposed to neuronal culture (feeder layer) before and after sustained depolarization plus pre-incubation with transcription inhibitor (DRB. Bar plot represents the mean. P value: ratio paired t-test. Error: SD (H) RNA quantification in primary astrocytes exposed to neuronal culture (feeder layer) beforeand after sustained depolarization (left) plus pre-incubation with transcription inhibitor (DRB) (right)(Nanostring nCounter, methods). Cdr1as and Cyrano shown in red. RNA counts are normalized to housekeeping genes (Actb, Tubb5 and Vinculin). Each dot represents the mean of 3 biological replicates (3 independent primary cultures from 3 animals). Grey line: linear regression. (I) RNA quantification of genes of interest (Nanostring nCounter, methods) in nuclear versus cytoplasmatic fractions of WT neurons DIV21. Each bar represents the mean difference of 3 independent biological replicates. Cdr1as and Cyrano: green. Primary miRNAs: red and orange. RNA counts are normalized to housekeeping genes (Actb, Tubb5 and Vinculin). (J) circRNA expression changes for WT neurons before and after K+ treatment. Plotted is the mean change of 2 independent biological replicates per condition. Red dot: Cdr1as. Blue and brown dots: other statistically significant circRNas (Methods).

**Supplementary Figure 2.**
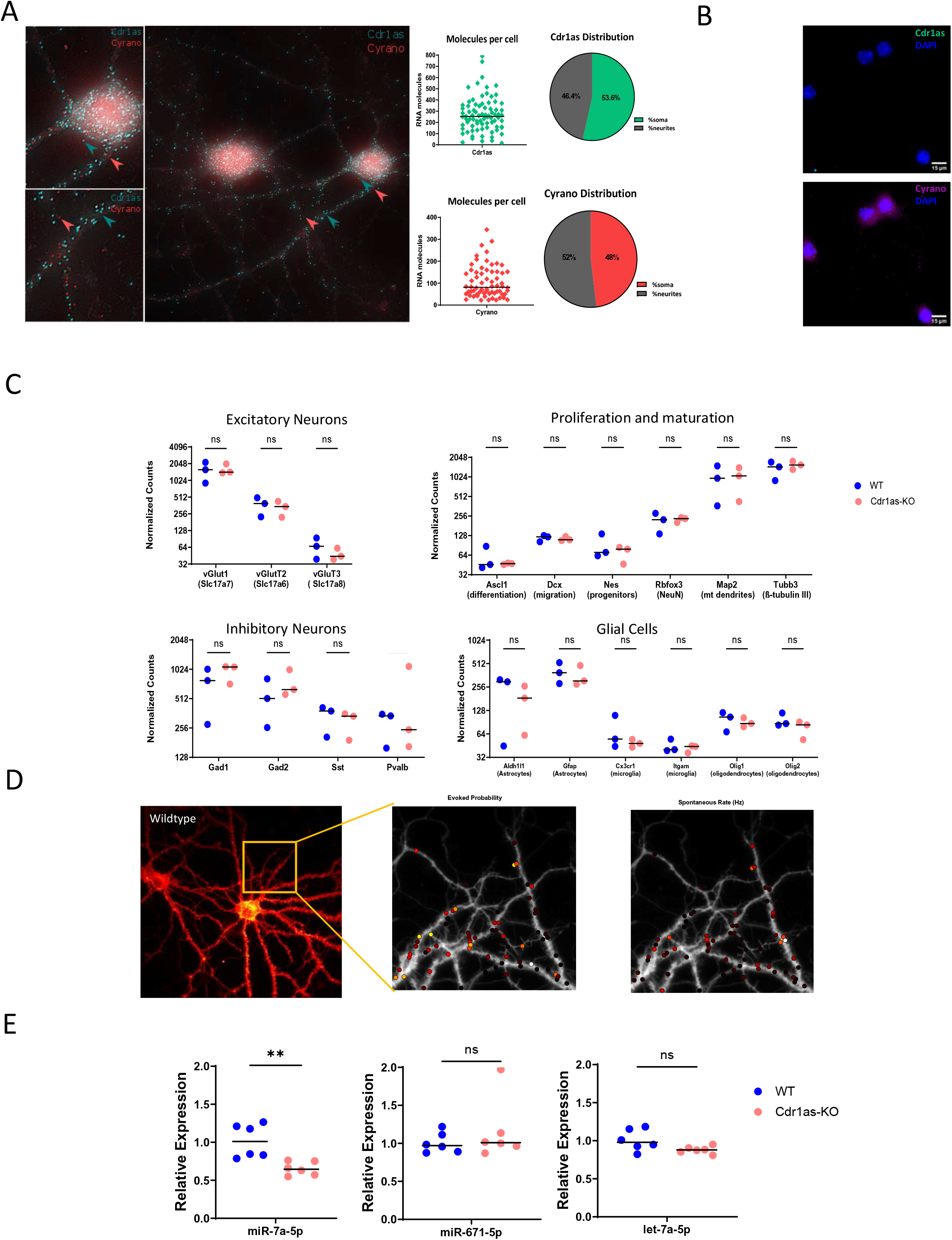
**(A)** Right: Single molecule RNA FISH (Stellaris, Methods) of Cdr1as (cyan) and Cyrano (magenta) performed in WT neurons. Left: smRNA FISH quantification of Cdr1as molecules (following Raj et al., 2008, methods). Each dot represents the mean number of molecules in an independent cell (soma + neurites) (Cdr1as n=6; 80 cells) (Cyrano n=5; 69 cells). Horizontal line: Median. Pie chart show molecule distribution in somas versus neurites. (B) Single molecule RNA FISH (Stellaris, Methods) of Cdr1as (cyan) and Cyrano (magenta) performed in Cdr1as-KO neurons DIV21. DAPI: blue **(C)** Quantification of cellular markers to characterize WT versus Cdr1as-KO primary cultures DIV21 (Nanostring nCounter, Methods). RNA counts are normalized to housekeeping genes (*Actb, Tubb5 and Vinculin*). Each dot represents an independent biological replicate (3 independent primary cultures from 3 animals). Excitatory neurons (*Sclc17a7, Sclc17a6, Sclc17a8*),Inhibitory neurons (*Gad1,Gad2,Sst,Pvalb*), Proliferation and maturation markers (*Ascl1,Dxc,Nes, Rbfox3, Map2,Tubb3*) and Glial cells markers (*Aldh1,Gfap, Cx3cr1, Itgam, Olig1, Olig2*) are plotted. P value: U Mann-Whitney test. Horizontal bar: Median. **(D)** Visualization of glutamate sensor expression (GlusnFR) in WT primary neuron DIV20 (representative image). Region of interest zoom in (yellow box). Middle and right panels indicate selection of active synaptic terminals and quantification of evoked probability and Spontaneous frequency, respectively. **(E)** Quantification of mature miR-7a-5p, mir-671-5p and let-7a-5p (TaqMan Assay, Methods) to characterize WT versus Cdr1as-KO primary cultures DIV21. RNA normalized to housekeeping genes (snRNA *U6,snoRNA 202*) Each dot represents an independent biological replicate (6 independent primary cultures from 6 animals).P value: U Mann Whitney test. Horizontal line: Median.

**Supplementary Figure 3.**
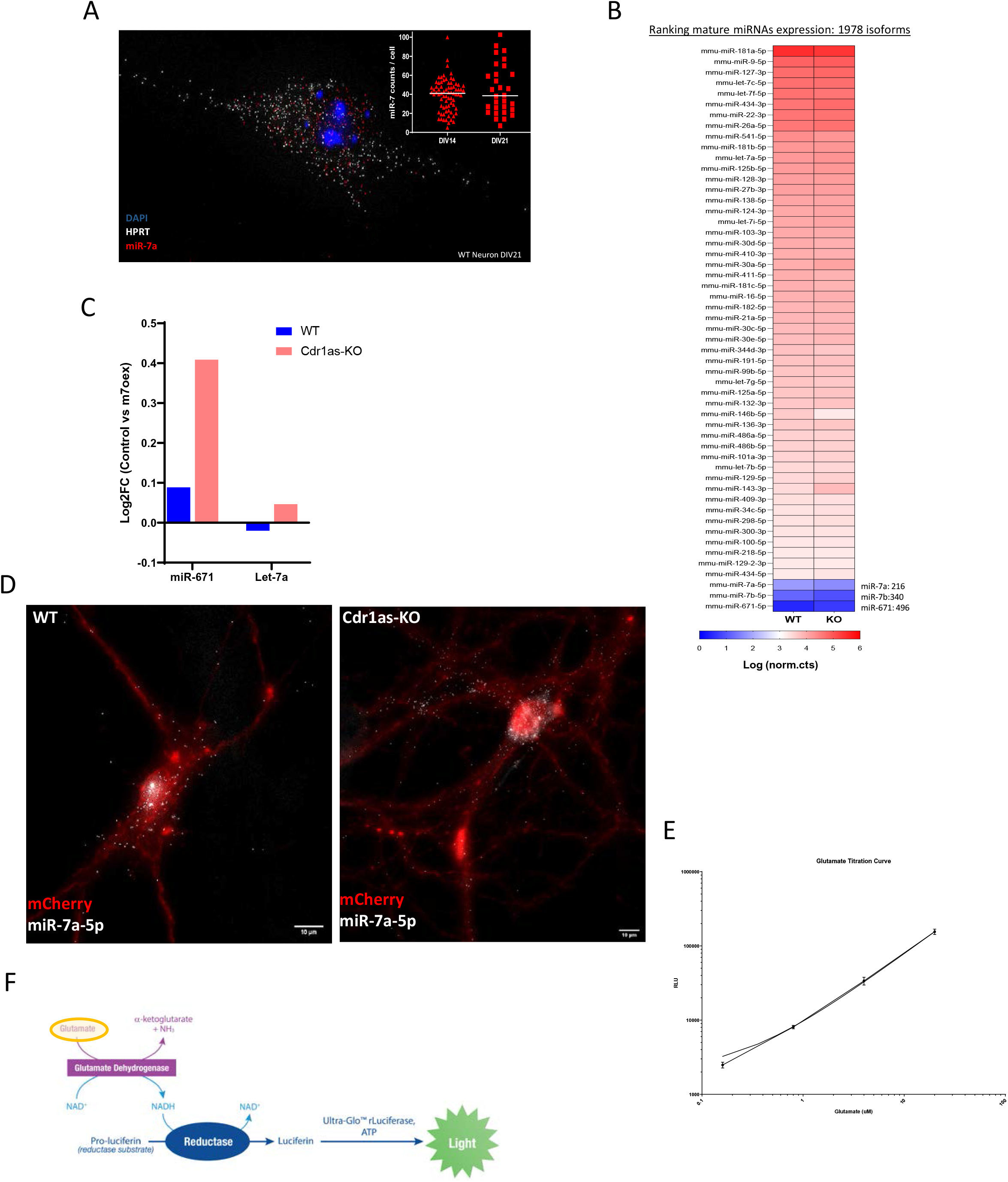
**(A)** Single molecule RNA FISH (ViewRNA Plus, Methods) of miR-7a-5p (red) and housekeeping gene *Hprt* (white) performed in WT neurons DIV14 and DIV21, DAPI: blue. Insert: smRNA FISH quantification of miR-7 molecules (following Raj et al., 2008, methods). Each dot represents the mean number of molecules in an independent cell: DIV14 (n=75 cells) and DIV21 (n=30 cells). **(B)** Heat map from bulk smRNA-Seq (Methods) (1978 miRNA isoforms detected), normalized expression plotted (log.norm.counts) of top 50 mature miRNAs in WT and Cdr1as-KO primary neurons DIV21, plus miR-7a, miR-7b and miR-671. **(C)** Quantification of miR-671 and let-7a after miR-7 overexpression (14dpi) by RNA-Seq (Methods) in WT and Cdr1as-KO neurons at DIV21. Bar plots represents mean of 4 independent biological replicates per genotype. **(D)** Single molecule miRNA *in situ* hybridization (ViewRNA Plus, Methods) of miR-7a-5p (white) and infection reporter mCherry (red), performed in WT and Cdr1as-KO neurons DIV21, 14 days post miR-7 overexpression (Widefield microscopy 60X) **(E)** Standard curve for calibration of glutamate secretion assay. Serial dilutions curve of 50uM Glutamate stock solution. Quantification of secreted glutamate concentrations for tested samples based on interpolation of RLU values. **(F)** Schematic representation of the enzymatic principle behind glutamate secretion assay. Modified from Glutamate-Glo™ Assay (Methods). GDH enzyme catalyzes the oxidation of glutamate with associated reduction of NAD+ to NADH. In the presence of NADH, Reductase enzymatically reduces a pro-luciferin to luciferin. Luciferin using Ultra-Glo™ Luciferase and ATP, and the amount of light produced (RLU) is proportional to the amount of glutamate in the sample.

**Supplementary Figure 4.**
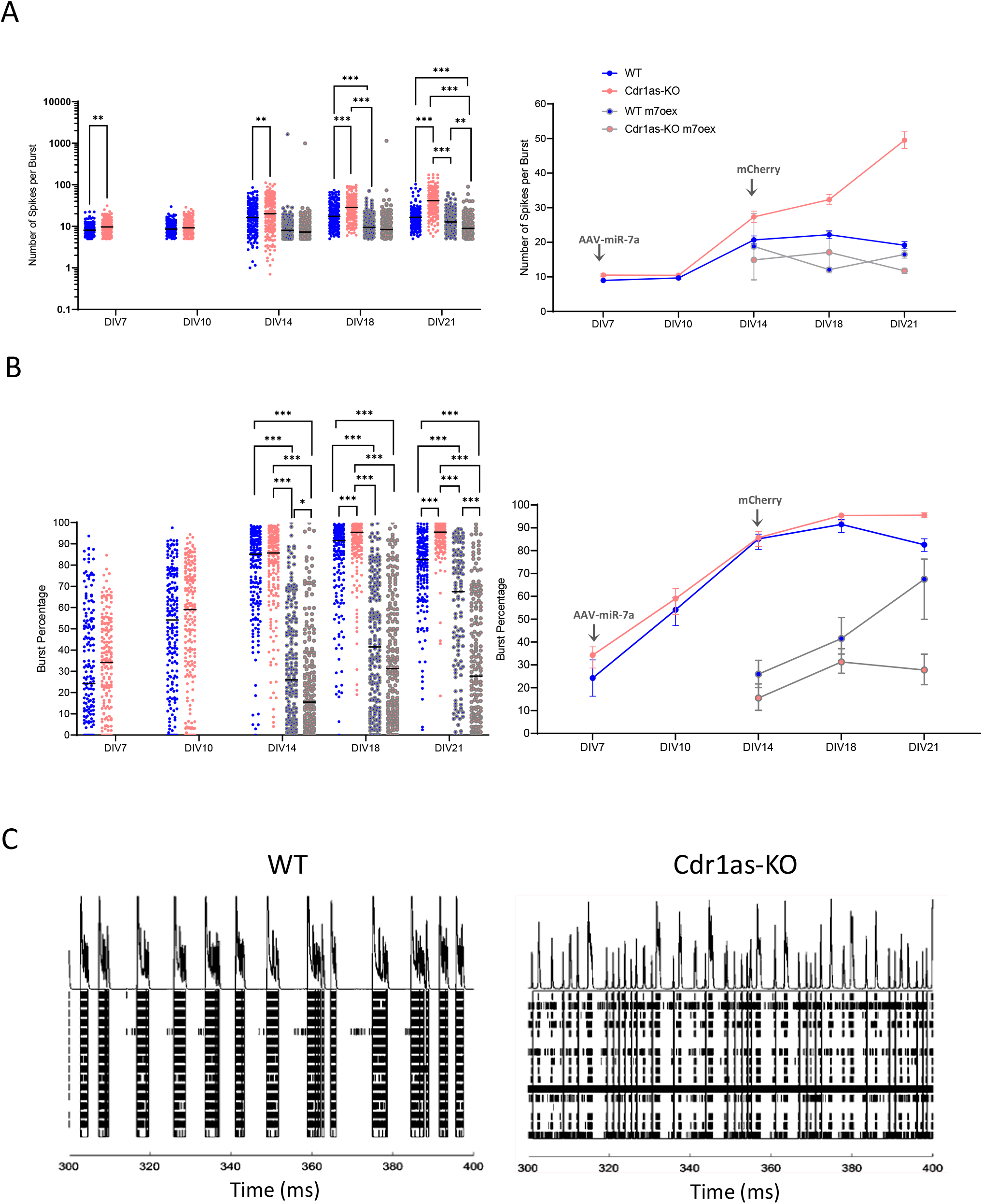
**(A)** Number of Spikes per Burst: Average number of spikes in a single-electrode burst. Left panel: each dot represents a single electrode recording from 4 independent primary cultures (WT = 187; WT + miR-7 overexpression = 194; Cdr1as-KO = 189; Cdr1as-KO + miR-7 overexpression = 200 electrodes). Horizontal line: median. Right panel: median value across timepoints with 95% confidence interval is plotted; arrows indicate transduction time (AAV, DIV7) and reporter first visualization (mCherry, DIV14), respectively. P value: two-way ANOVA, all other comparisons were non-statistically significant (not shown). **(B)** Burst Percentage: number of spikes in single-electrode bursts divided by the total number of spikes, multiplied by 100. Left and right panels plotted as in (A). **(C)** Example raw spikes from multi electrodes Array recording: 100 ms raw spikes of WT and Cdr1as-KO neurons DIV21. Each row represents one independent electrode (blac bars).

**Supplementary Figure 5.**
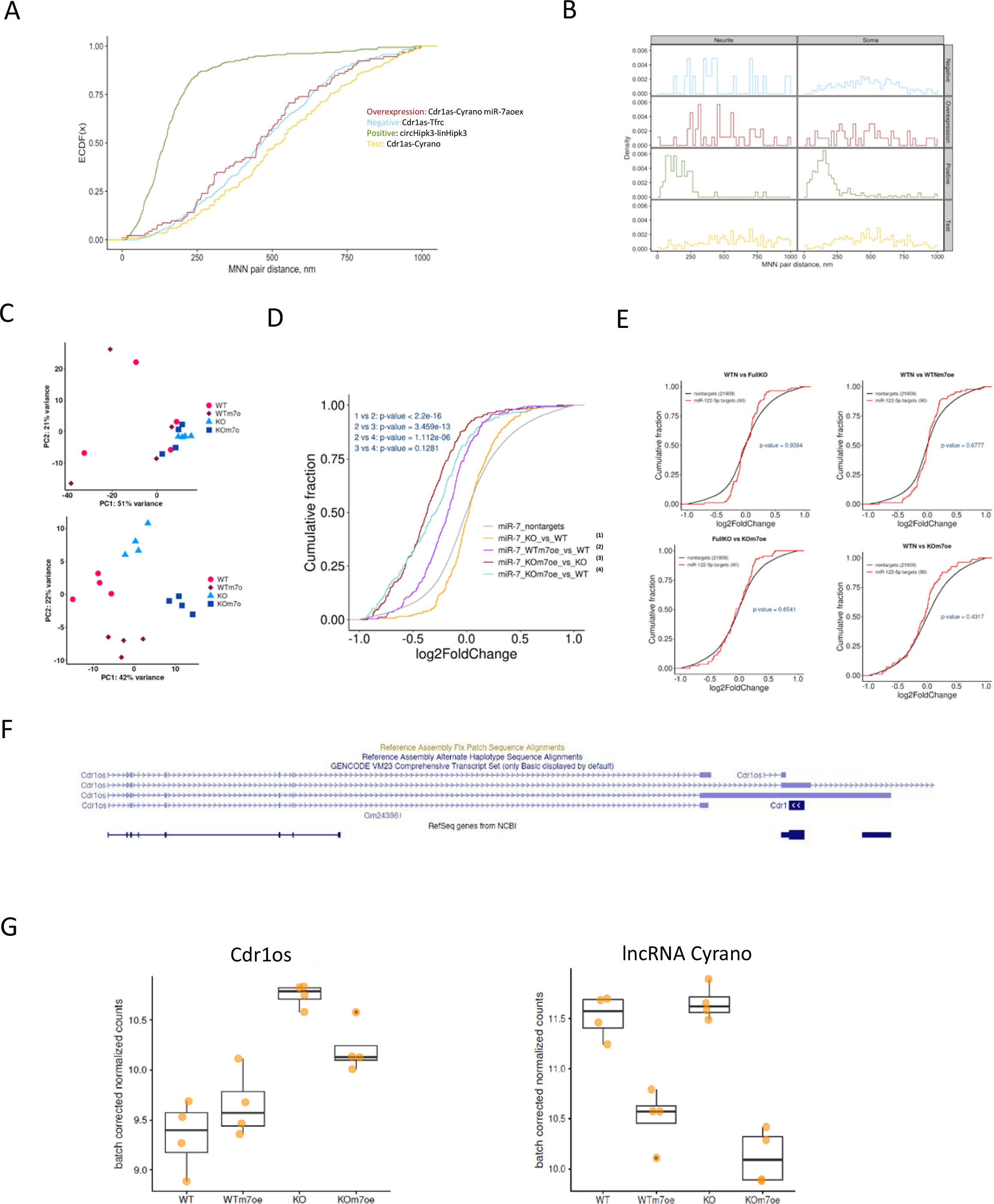
**(A)**Cumulative distribution function (CDF) plot of molecule distances based on smRNA FISH images of WT neurons DIV21. before and after miR-7 overexpression (Methods) comparing all computationally predicted mutual nearest neighbors distances in nm all conditions (MNN, Methods). Positive technical control: circHipk3-linHipk3; Negative control: Cdr1as-Tfrc; Test: Cdr1as-Cyrano; overexpression: Cdr1as-Cyrano miR-7aoex. 2 independent biological replicates from 2 animals. **(B)** Density plot molecule distances based on smRNA FISH to compare somas versus neurites for all tested conditions. Analysis conditions same as in (A). **(C)** Principal component analysis (PCA) of all data sets. Each replicate represented by one dot. Original (up) and batch corrected with nested design (down). **(D)** Cumulative distribution function (CDF) plot of gene expression comparing all computationally predicted miR-7 targets (Methods) to mRNAs lacking predicted miR-7 target sites (non-targets, grey), across 4 independent biological replicates of WT, Cdr1as-KO, WT + miR-7 overexpression, Cdr1as-KO + miR-7 overexpression. P value: U Mann–Whitney test. **(E)** Cumulative distribution function (CDF) plot of gene expression comparing all computationally predicted miR-122 targets (Methods, red) to mRNAs lacking predicted miR-7 target sites (non-targets, black), across 4 independent biological replicates of WT, Cdr1as-KO, WT + miR-7 overexpression, Cdr1as-KO + miR-7 overexpression. P value: U Mann–Whitney test. **(F)** Cdr1os transcript (Cdr1as precursor transcriptional unit)reference sequence alignments from Genome Browser (GENCODE VM23) **(G)** Gene expression of Cdr1os and lncRNA Cyrano in each data set. Box plots, 4 independent biological replicates per condition, for each comparison tested. (FDR < 0.05).

**Supplementary Figure 6.**
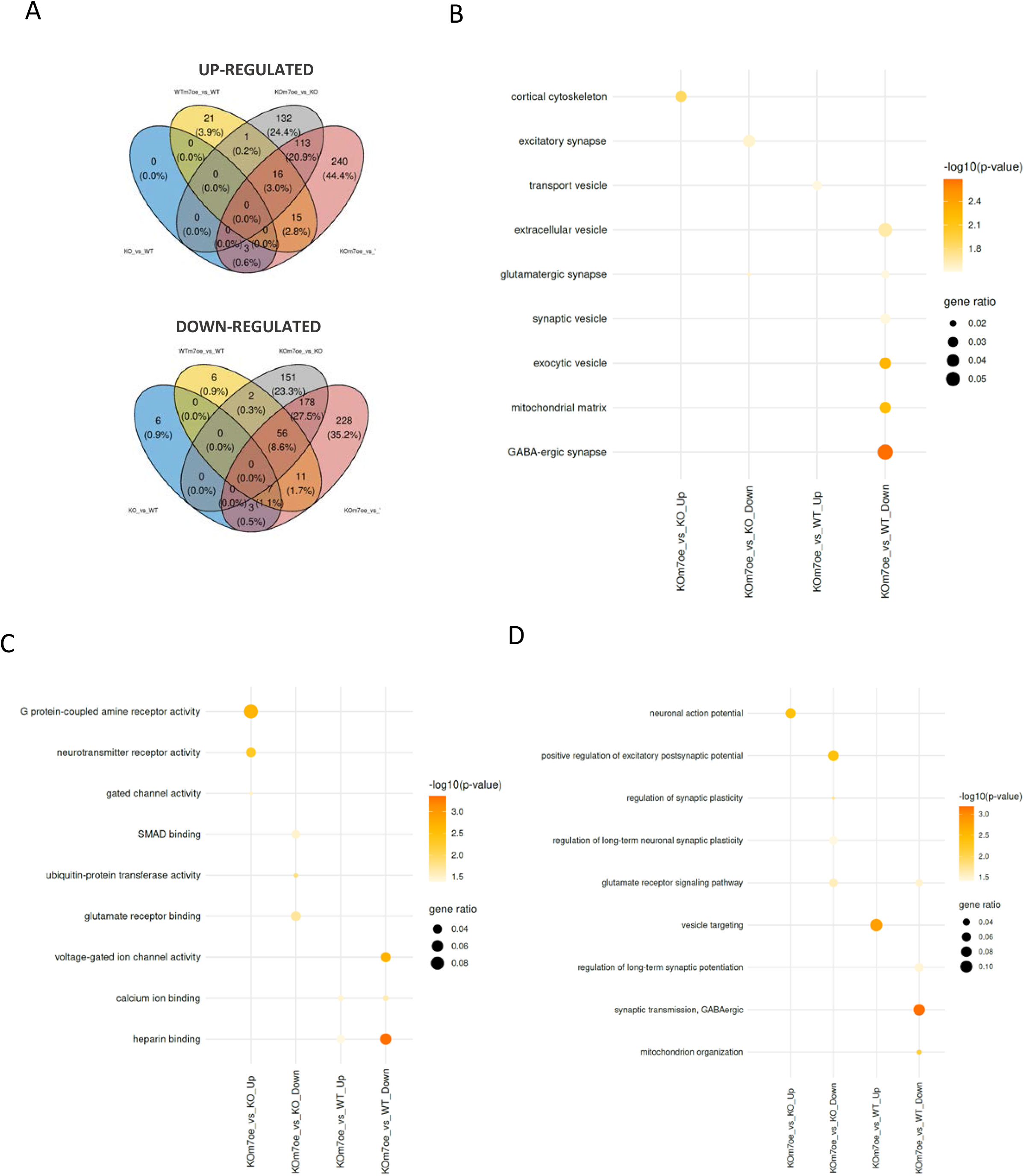
**(A)** Number of specific and commonly up-regulated and down-regulated genes from mRNA expression changes for each comparison tested (Cdr1as-KO vs WT, WTmiR-7oex. vs WT, Cdr1as-KO miR-7oex.vs Cdr1as-KO). Venn diagram, mean change of 4 independent biological replicates per condition. **(B)** Enrichment profile from gene ontology (GO) analysis of cellular compartments (CC). Gene ontology enrichment analysis done using topGOtable function in the pcaExplorer (Marini et al., 2019, Methods). The dot size shows gene ratio and the color denotes the FDR-corrected p-value. Genes with average log2 fold change of 0.5 and adjusted p-value less than 0.05 were considered significant and all expressed genes were used as background. **(C)** Enrichment profile from gene ontology (GO) analysis of Molecular Functions (ML). Enrichment profile from GO analysis plotted as in (B) **(D)**Enrichment profile from gene ontology (GO) analysis of Biological Process (BP). Enrichment profile from GO analysis plotted as in (B)

**Supplementary Figure 7.**
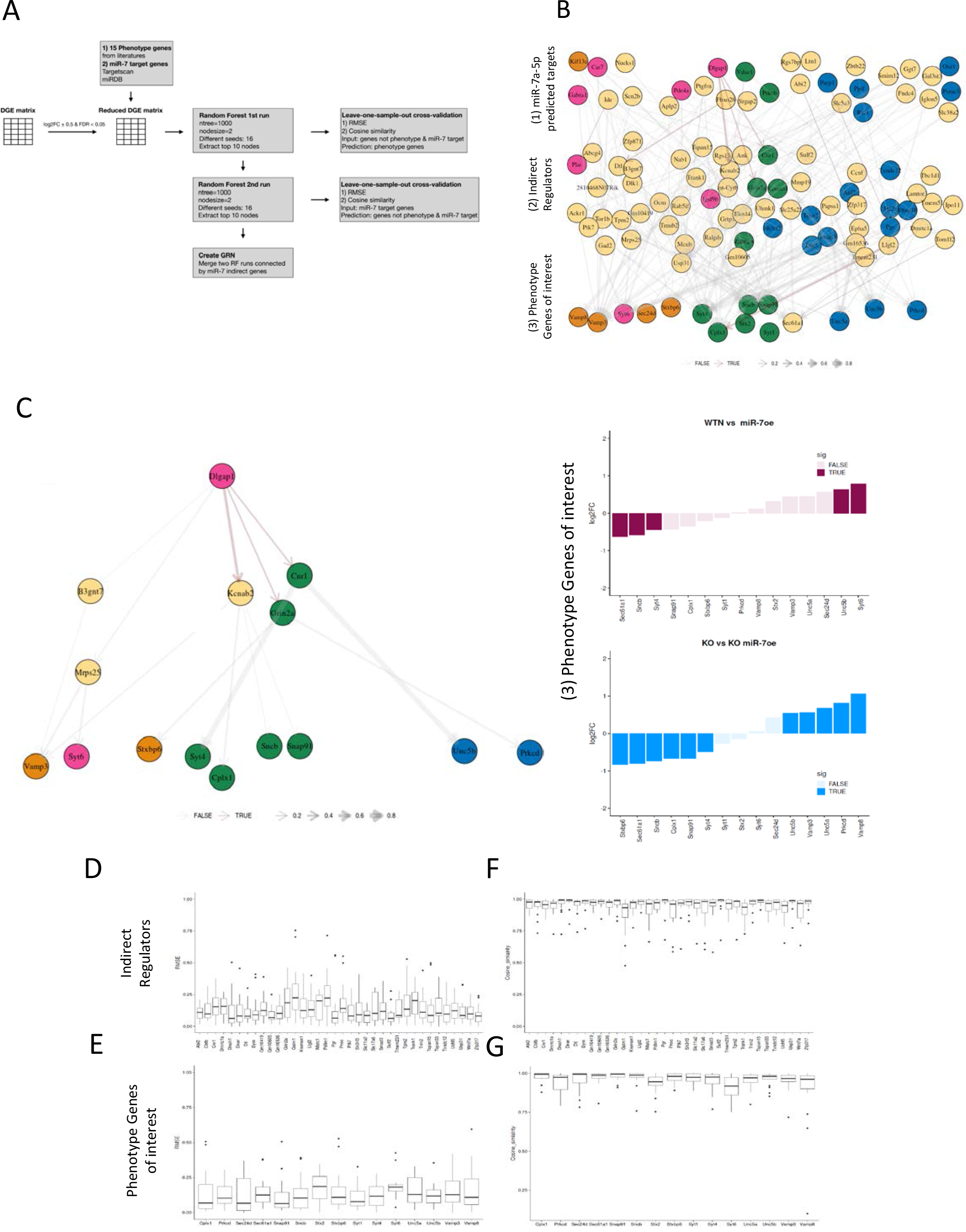
**(A)** Schematic representation of Random Forest modelling methodology. **(B)** Consensus gene regulatory network (GRN) of miR-7 and Cdr1as in neurons. The first layer are miR-7 target genes, the middle layer are intermediate regulator genes and the bottom layer are phenotype-related genes from literature. The node color indicates gene ontology terms associated with synaptic (pink), vesicle (orange), apoptosis (blue), and common (green), respectively. The edge width corresponds to the relative fraction of cross-validation models having this edge (Methods). Colored edges indicate links found in StringDB (Methods). **(C)** Left: Selected connection of the inferred GRN with the largest number of recovered known links from StringDB. Right: Differentially expressed genes in phenotype-related genes of GRN network in each comparison. Bar plots, the mean log2 fold change of 4 independent biological replicates per condition, for each comparison tested. Transparent bars, not significant changes (FDR > 0.05). **(D)** Leave-one-out cross-validation performance for “indirect regulators” layer. For each gene, one sample out of 14 was left out and RMSE score was calculated for each of the remaining 13 samples (Methods). **(E)** Leave-one-out cross-validation performance for “Phenotype genes of interest” layer. For each gene, RMSE score was calculated as in (D). **(F)** Leave-one-out cross-validation performance for “indirect regulators” layer. For each gene, one sample out of 14 was left out and cosine similarity score was calculated for each of the remaining 13 samples (Methods). **(G)** Leave-one-out cross-validation performance for “Phenotype genes of interest” layer. For each gene, cosine similarity is calculated as in (F)

## Supplementary Discussion

Neuronal activity regulated secretion pathways are the key mechanistic tools to control the network connectivity responsible for long-term brain changes (LTP, synaptic morphological changes, neural coding, etc.) and subsequent plastic adaptations (learning, memory, conditioning), where transcriptional and post-transcriptional changes are necessary for the development of neuroplasticity, even though transcriptional target genes that are involved in long-term plasticity have yet to be identified (McClung & Nestler, 2008). Therefore, the importance of this ncRNA regulatory network in an *in vivo* biological context remains to be tested. We believe that finding scenarios where cortical neuronal networks need to adapt quickly and permanently to stressed or highly triggered conditions would be the perfect context to further examine the mechanistic roles of the Cdr1as and miR-7 axis.

According to the description done by Rybak-Wolf et al. (2015) in primary hippocampal neurons Cdr1as expression increases dramatically on DIV14 (120 FC), which could indicate that the role of Cdr1as in forebrain is relevant for neuronal function in mature states of synaptic connections. This positively correlate with our observation of increasing neuronal activity over time in Cdr1as-KO neurons (**Fig. 3 and 4)**, where the action of miR-7 must be more strongly buffered.

The precise intracellular pathways activated to achieve Cdr1as up-regulation after stimulation and all observed neuronal responses dependent of miR-7 and Cdr1as regulations remain to be uncovered. Some indications of induction of Cdr1as expression by several secretagogues that activate cAMP and PKC signaling pathways (Xu et al., 2015), which in neurons are related to the activation of metabotropic glutamate receptors (Niciu et al., 2012), combined with our gene ontology enrichment analysis which indicate that vesicle targeting and glutamate receptors signaling pathways are affected only in Cdr1as-KO neurons after miR-7 sustained up-regulation (**Fig. 6D**).

A small dysregulation of extremely low expressed miR-7, due to the constitutive loss of Cdr1as (**Suppl. Fig. 2D and 3B**) is most likely not reflected in global abundance of mRNAs, because miRNAs are fine-tuners of transcriptome regulations that mostly act under time and stimulus specific circumstances (Bartel, 2004; Schratt, 2009), therefore, only the perturbation of low abundant miR-7 allows us to observe global RNA changes.

Unexpected large indirect up-regulation of non-miR-7a-targets genes, which can be related with the compensatory mechanism activated by mir-7 to compensate dysfunctional glutamate secretion of Cdr1as-KO neurons.

Cdr1os (Cdr1as precursor; **Suppl. Fig. 5F**), which previously was shown to be up-regulated by the loss of Cdr1as locus in bulk brain tissue (Barrett et al., 2017), is not altered by sustained miR-7 overexpression (**Suppl. Fig. 5G, left panel**), an RNA sequencing also confirmed the significant down-regulation of Cyrano in both genotypes caused by miR-7 up-regulation, but not affected by the loss of Cdr1as itself. (**Suppl. Fig. 5G**, right panel).

All together in this research work we discovered that sustained miR-7 overexpression in Cdr1as-KO neurons specifically regulates set of genes of functional pathways related to synaptic signaling dynamics, equilibrium of excitatory-inhibitory transmission and synaptic plasticity, and those gene pathway regulations are reflected in modulations of excitatory neurotransmitter release, local and network synaptic activity of the neurons.

Furthermore, we proposed potential key gene regulatory pathways connecting gene ontology terms with our functional phenotypes. To identify genes associated with the regulation of neuronal phenotypes of sustained miR-7 expression in neurons with the loss of Cdr1as, we performed Random Forest (RF) modelling (**Suppl. Fig. 7A**), which yielded in 154 candidate genes across three regulatory layers: (1) miR-7a-5p predicted targets; (2) ‘intermediate regulator’genes; (3) phenotype-related genes of interest (**Suppl. Fig. 7B**). Then, we used a known protein-protein interaction network (StringDB) to select reliable connections in the predicted gene regulatory network (red arrows in Suppl. Fig. 7B).

Out of the predicted miR-7a-5p target genes, Dlgap1, a postsynaptic scaffold protein in neuronal cells, had the most reliable connections (**Suppl. Fig. 7C**). Dlgap1 regulates directly the Kcnab2, Grin2a and Cnr1 genes, which encode for a potassium channel subunit, a glutamate receptor and a G protein-coupled cannabinoid receptor, respectively. These genes have predicted connection to the following phenotype-related genes: Cplx1, Snap91, Sncb, Vamp3, Prkcd, Stxbp6, which directly induce changes in neurotransmitter secretion. Some of these genes have been previously linked to miR-7-dependent secretion phenotypes (LaPierre et al., 2022; Latreille et al., 2014; Xu et al., 2015).

